# Robust, Fiducial-Free Drift Correction for Super-resolution Imaging

**DOI:** 10.1101/2021.03.26.437196

**Authors:** Michael J. Wester, David J. Schodt, Hanieh Mazloom-Farsibaf, Mohamadreza Fazel, Sandeep Pallikkuth, Keith A. Lidke

**Affiliations:** Department of Mathematics and Statistics, University of New Mexico, Albuquerque, NM 87131 USA; Department of Physics and Astronomy, University of New Mexico, Albuquerque, NM 87131 USA; Lyda Hill Department of Bioinformatics, UT Southwestern Medical Center, Dallas, TX 75390 USA; Center for Biological Physics, Department of Physics, Arizona State University, Tempe, AZ 85287 USA

## Abstract

We describe a robust, fiducial-free method of drift correction for use in single molecule localization-based super-resolution methods. The method combines periodic 3D registration of the sample using brightfield images with a fast post-processing algorithm that corrects residual registration errors and drift between registration events. The method is robust to low numbers of collected localizations, requires no specialized hardware, and provides stability and drift correction for an indefinite time period.

## Introduction

Drift correction is a prevalent problem in microscopy, in which the sample being examined alters its position over time, leading to distortions for data collected over a series of movie frames. Typically, translational motion is the main significant positional change, which is what we will consider here, however, others have also examined the issue of rotational movements^1,2^. Drift can be due to a variety of factors, such as temperature variation, vibration and mechanical relaxation of the measuring instruments^3–5^, becoming significant for long recording times.

Drift occurs in all varieties of microscopy, for example, scanning electron microscopy (SEM)^1,6^, scanning (electron) tunneling microscopy (STM)^7,8^, and scanning probe microscopy^9^ among others. Here, we are concerned with super-resolution microscopy of blinking fluorescent particles in which the images are taken over many movie frames^10–13^.

Various techniques to eliminate drift errors have been developed, such as producing specialized hardware and introducing fixed fiduciary markers or patterns as reference points^14–16^. Some researchers have suggested tracking image features, for example, the center of mass of a cell-like object^17^. Several techniques involve identifying and extracting image features, then matching them for global correspondence^6^, using clusters of those features^2^, or their nearest (feature) neighbors^8^ in order to deduce the drift. These techniques work best with non-pointlike features^6^.

Active drift correction via continuous brightfield / non-fluorescent imaging with feedback to the stage position has been used by some^18,19^, the latter using circular intracellular vesicles to substitute for fiducial markers. Tang et al.^4^ minimized the normalized root-mean-square error between brightfield images over all pixels.

A common technique for estimating drift post experiment is by using multiple localizations from the same source that are interspersed throughout the data collection. The most common drift correction technique for super-resolution images using point localization data has been image cross-correlation, in particular, a fast implementation using fast Fourier transforms (FFTs) known as phase correlation^20–22^. A variation of this technique involves auto-correlation at time zero and cross-correlation subsequently^3^. McGorty et al.^23^ used cross-correlation for 3D drift estimation.

Two additional extensions involve sum images, in which the sum of a series of frames is used to more reliably indicate localizations. In Wang et al.^24^, each frame is assumed to contain an incomplete description of an identical structure. Groups of frames are combined into sum images. Redundant cross-correlation is then used, considering all possible combinations of the sum images. The second procedure^25^ identifies the FFT cross-correlation maximum between a sequential pair of frames. The estimated drift vector is applied to the later frame, which is then added to the sum image of the earlier shifted frames (or simply earlier frame on the first iteration). The process then repeats with the earlier frame now being a sum image.

Two techniques directly employing point positions are using a Bayesian statistical framework to calculate the drift corrections for every image frame^26^ or per emitter when grouping localizations^27^. A third technique involves maximizing the molecular constraint field (MCF) cost function, where the MCF is a function of the distances between the points in two images^5^. The MCF cost function, defined as the sum of the negative exponentials of the distances between all points in a fixed reference image with respect to all points in a shifted movable image, takes into account the sparseness of points around each position, and produces near zero contribution when the distances become large. Typical usage involves combining collections of sequential frames into datasets (i.e., sum images) to which the MCF is applied incrementally while combining previously drift corrected results with the reference image.

Here, we propose a combination of brightfield registration performed during an experiment before each collection of super-resolution frames forming a dataset, followed by a post procedure that uses the nearest neighbor distances of the localizations found to perform drift correction. All software described was written in MATLAB.

## Algorithm

### Brightfield registration

For brightfield registration, we establish a protocol in which super-resolution imaging is performed for a sequence of movie frames, which we term a dataset. A *z*-stack of brightfield reference images is obtained before data collection begins. Before each dataset is collected, a new brightfield *z*-stack is obtained and compared with the previous reference *z*-stack, finding the best *z* and (*x, y*) fits via a scaled cross-correlation, allowing the reference *z*-stack and the current *z*-stack to be aligned. The procedure then repeats with the next dataset. See Fig. 1(a). A complete data collection for one sample typically includes 10-50 datasets.

**Figure 1.**
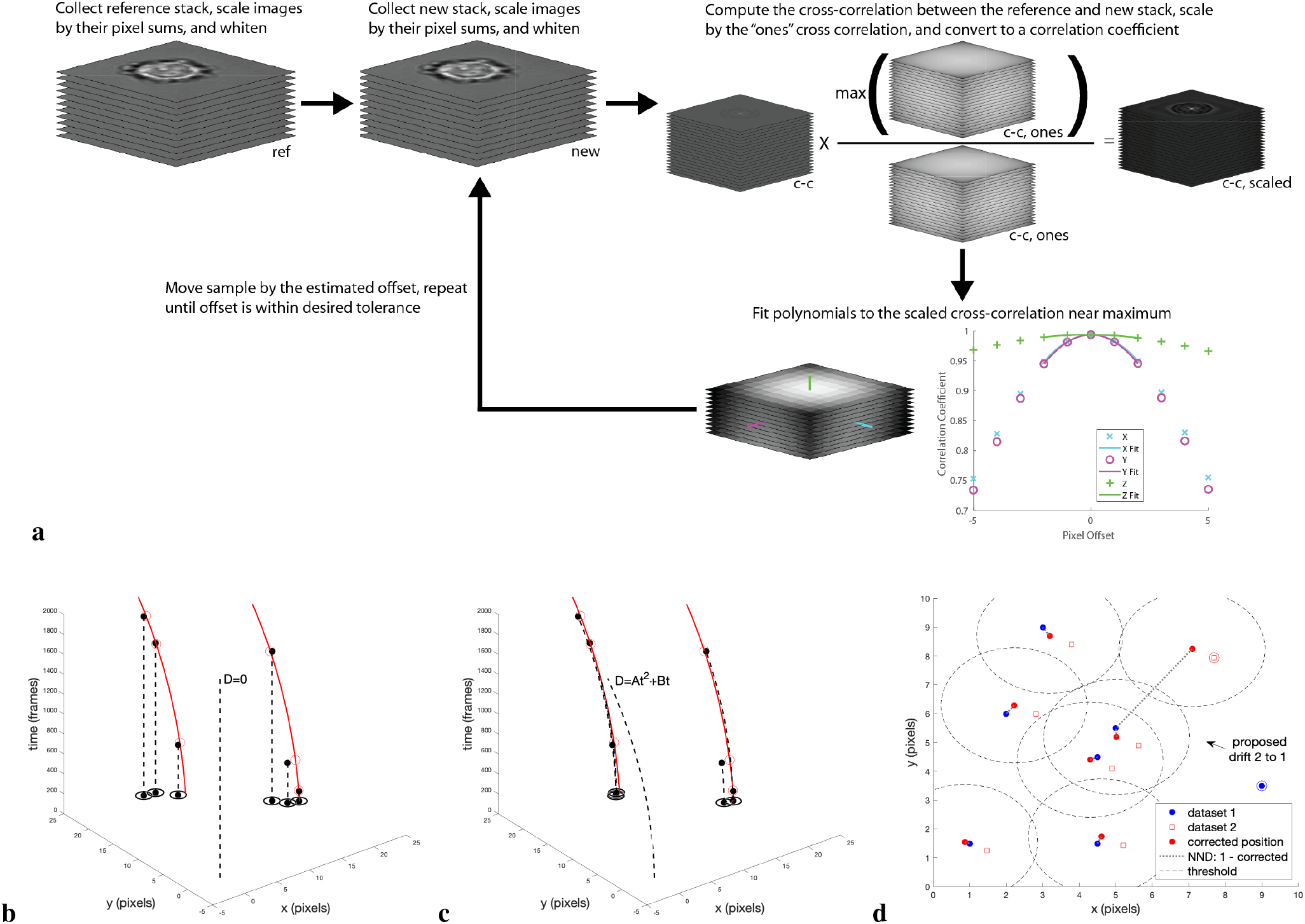
Concept of (a) brightfield registration and examples of (b,c) intra-dataset and (d) inter-dataset post-processing drift correction scenarios. (b,c) The red lines represent the true drift curves for two emitters (red circles) at different time points. The black dots are the nearby emitters’ observed localizations, which are projected by a particular model into the *x − y* plane via the dashed lines. (b) models no drift, while (c) models quadratic drift. The isolated dashed lines demonstrate the model used. The black circles represent the intra-dataset nearest neighbor distance threshold around instances of the *x − y* projected localizations. (d) The blue dots represent dataset 1 localizations. The hollow red squares are dataset 2 localizations. The red dots are predictions from a drift model where the dataset 2 localizations are assumed to have undergone a lateral shift and now their original locations in dataset 1 are estimated from the drift model. Note that the corrected positions are offset from the dataset 2 localizations by the constant vector expressed by the arrow on the right. The ringed symbols on the right side of the plot represent emitters that are on in only one dataset (which one indicated by their color). The dotted gray lines connect the nearest dataset 1 neighbor of each drift model corrected position. The dashed circles represent the inter-dataset nearest neighbor threshold around the drift model corrected positions. Note that the nearest neighbor of the drift model corrected position of the upper right highlighted dataset 2 localization exceeds the threshold.

For a single dataset, registration between the reference *z*-stack, ref, and the new brightfield *z*-stack, new, proceeds as follows. Images in each of the two stacks are individually scaled by the sum of their pixel values:

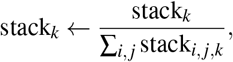

where stack_*k*_ denotes a single image taken from either ref or new and stack_*i,j,k*_ denotes a single pixel in stack_*k*_. This process is done for every image in both ref and new. Scaling each image in this manner reduces biases in the *z* location of the cross-correlation maximum that may occur due to differing intensities of images taken at different *z* positions. The two stacks are then individually whitened:

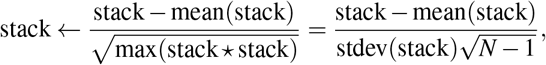

where ✶ denotes cross-correlation, *N* is the total number of pixels in the *z*-stack, and mean/stdev(stack) is the mean or standard deviation of pixel values across the entire *z*-stack. The whitening procedure will both reduce biases in the location of the cross-correlation maximum (i.e., for image stacks with a non-zero mean pixel value, the cross-correlation maximum is biased to correspond to small offsets between the stacks relative to the size of the stacks) and will scale the stacks such that pixel values in the final cross-correlation stack will range from −1 to 1. A scaled 3D cross-correlation between ref and new is then computed as

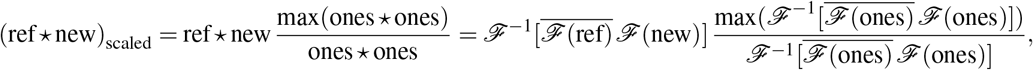

where 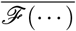 indicates the complex conjugate of a fast Fourier transform (denoted by 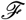) and ones denotes a stack of 1’s the same size as the stacks ref and new. Each pixel in ones ✶ ones will contain a count of the total number of overlapping pixels between ref and new that were used to compute each pixel in ref ✶ new. The element-wise scaling of ref ✶ new by ones ✶ ones will reduce the bias in the location of the cross-correlation maximum introduced by the differing number of overlapping pixels used to compute each pixel in ref ✶ new. The drift is then independently computed for each of *x, y* and *z* by fitting second-order polynomials to the cross-correlation values along lines of pixels in the dimension being fit which intersect the maximum cross-correlation pixel in the 3D volume.

### Post-processing drift correction

Once all the datasets are collected, the post-processing drift correction procedure is applied to the data. This procedure divides the problem into intra-dataset and inter-dataset drift correction, both minimizing a cost function that is simply the thresholded sum of the nearest neighbor distances for all the predicted emitter positions in a dataset, either with respect to itself for intra-dataset drift correction (noting that different sets of localizations over time are coming from the blinking fluorophores), or the first dataset for inter-dataset drift correction. Intra-dataset drift correction is performed first, with the results saved for the next phase. For inter-dataset drift correction, the datasets are drift corrected in sequence against the first dataset. Note that the inter-dataset drift correction is really a dataset registration process. Figure 1(b,c,d) illustrates the basic concepts of these two procedures.

### *k*-nearest neighbor search

The core of these processes involves *k*-nearest neighbor searches in which the image is partitioned by a data structure called a *k-d* or *k*-dimensional tree. Each step in the *k-d* algorithm splits a region of the image space by a hyperplane, dividing the points into one of two half-spaces. The regions are formed by earlier splits. *k-d* trees are typically constructed by cycling through the dimensions (*x* then *y* then *x* then … in 2D) as regions are split further and further, and choosing a median splitting plane that evenly divides the points between the two half-spaces. In this situation, the *k-d* tree is considered balanced. Nearest neighbor distances are quickly computed once a *k-d* data structure is produced. MATLAB implements balanced *k-d* trees for nearest neighbor searches^28^.

### Inter-dataset drift correction

To understand what is happening in more detail, consider Fig. 1(d), depicting the situation for inter-dataset drift correction (in 2D), which is easiest to explain first. The emitters in dataset 1 are assumed to undergo a lateral shift, that may occur between datasets because of small residual errors in the brightfield registration process, in the transition to dataset 2. Brightfield registration is performed in the interval, and it is assumed that the intra-dataset drifts have been removed during application of the first part of the post-processing drift correction algorithm. What is left then is simply a constant offset between dataset 1 and dataset 2 due to residual errors from the brightfield registration, actual sample drift during the typically short time interval, and other errors due to the dynamic behavior of the system. For each localization in dataset 1, the nearest neighbor in dataset 2 of the predicted location of the localization under assumptions of a lateral shift can be determined. For the sake of efficiency (see **Methods; Post-processing drift correction**), the model we actually use is the predicted location of dataset 2 localizations when unshifted to dataset 1. A sum of all the thresholded nearest neighbor distances (indicated by the connecting dotted lines) between the model (derived from dataset 2 and an assumed lateral shift) and the actual dataset 1 localizations is then constructed, which will become the cost function for an optimization routine. The minimum value of this sum, corresponding to the best correspondence between the model predictions and dataset 1 is searched for by the optimizer, yielding a model of lateral x- and *y*-shifts (and *z*-shifts in 3D) between these two datasets. The process then repeats using dataset 1 and dataset 3, etc.

### Intra-dataset drift correction

The intra-dataset algorithm, depicted in Fig. 1(b,c), is similar in many ways to the inter-dataset procedure. Here, the localizations, instances of true emitters, in all the frames in a single dataset are related. The drift model now assumes a polynomial dependence on time, so given the model parameters, each localization in the entire dataset is projected to *t* = 0 using the motion model. The constant in the polynomial fit is set to zero in order to match the model with the initial locations in the dataset. The first figure shows the situation in the case of no drift model, while the second provides an example using a quadratic time model. Once again, the nearest neighbor of each shifted localization in the entire dataset is determined, but this time with respect to the same model dataset (here, nearest neighbor means a different localization than the original one). Thus, minimizing the sum of the thresholded nearest neighbor distances minimizes the dispersion of the localizations throughout the dataset. In other words, we are trying to get the localizations to move as a group between frames and not spread out as would typically happen for poor drift models. This then carries over to the true emitters.

### Thresholding and initial guesses for the optimizer

The threshold for the nearest neighbor sums is chosen to de-emphasize nearest neighbors not likely to have been generated by the same emitter > *l_intra_* (= 1 pixel) away in the intra-dataset analysis, and > *l_inter_* (= 2 pixels) away in the inter-dataset portion of the algorithm. Mathematically, if *d_i_* is the nearest neighbor distance to the estimated localization *i*, the thresholded nearest neighbor sum is then

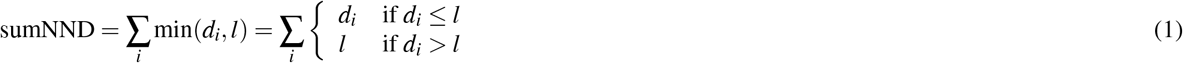

where *l* = *l_intra_* or *l_inter_* is the threshold. For intra-dataset movement, fluorophores are constantly blinking on and off and are also sparsely represented, so this threshold tries to prevent bad neighbor pairings with false large contributions to the nearest neighbor sum. For example, when the nearest neighbor of a fluorophore blinks off in one frame of a dataset, this can result in a totally different and potentially distant nearest neighbor for the same fluorophore in the next frame. If distant, this can produce a huge contribution to the nearest neighbor sum if no threshold was present. In the inter-dataset drift procedure, this is less of a problem because the data is much denser, however, a threshold can be important for sparse datasets (see Supplementary Fig. S4). See also Fig. 1(d) for an example of a threshold coming into play when an emitter only present in dataset 2 (red ringed square) has a predicted dataset 1 position that is far away from any actual dataset 1 localizations. For intra-dataset drift correction, an initial guess of zero for all coordinates is used for the optimizer; for inter-dataset drift correction, an initial guess based on the accumulation of the drift of those datasets that have been corrected in previous iterations is used (or zero if brightfield registration has been employed between datasets). The polynomial fits for drift correction are a function of frame (or dataset) number; in other words, time. By default, we use a linear fit intra-dataset and a constant fit inter-dataset, but these can be easily changed if desired.

This procedure runs in both 2D and 3D. See the **Methods** for additional details.

## Results

The drift correction procedure consists of two phases: brightfield registration before the acquisition of each dataset, and a post-processing drift correction algorithm (driftCorrectKNN).

### Application to 2D imaging

Figure 2 shows an example of the effects of not performing versus performing brightfield registration for alpha tubulin in HeLa cells using 2,000 frame datasets. Only brightfield registration took place; the post-processing drift correction algorithm was not applied here. In general, brightfield registration is not this dramatic, but for our day-to-day usage, frequently our post-processing drift correction algorithm has little work to do because the brightfield registration works so well.

**Figure 2.**
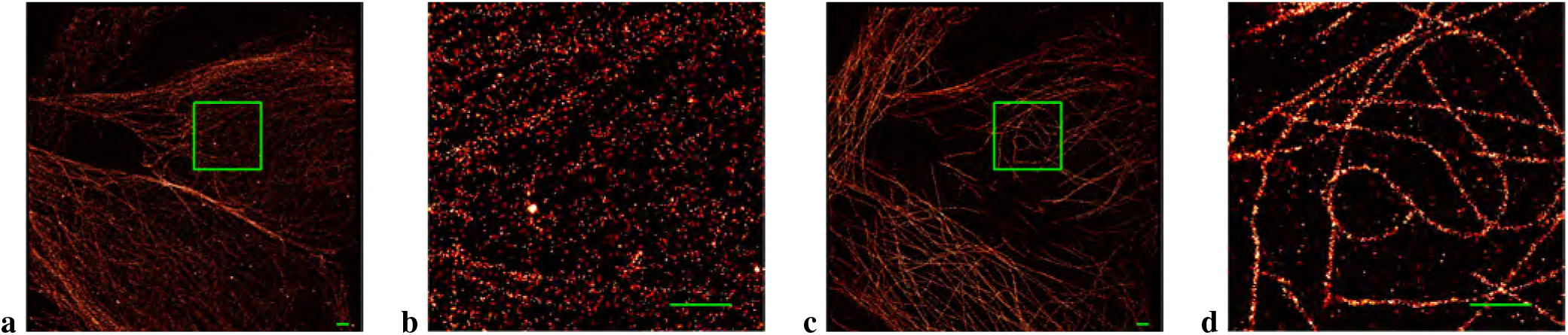
The effect of performing only brightfield registration. (No post-processing drift correction was employed here). (a) Alpha tubulin in HeLa cells imaged without employing brightfield registration between datasets. (c) The same cell imaged with brightfield registration performed before the acquisition of each dataset. 6 datasets containing 2,000 frames apiece were collected. (b,d) Zoomed in view of the selected region in the image to the left. All scale bars measure 1 *μ*m.

Figure 3 shows the results of applying our post-processing drift correction algorithm to two different 2D examples, one involving nanorods labeled via DNA-PAINT in which brightfield registration was not used, and the other consisting of actin microfilaments in HeLa cells labeled with fluorescent particles which did use registration. The first image in each sequence is the raw result from localization identification to which frame connection has been applied to consolidate emitters (see **Supplementary Methods**). The second image is the result of applying the drift correction algorithm. In each example, a small selected area of the entire region of interest (ROI) is also displayed. In the two examples, the drift correction algorithm considerably sharpens the observed details.

**Figure 3.**
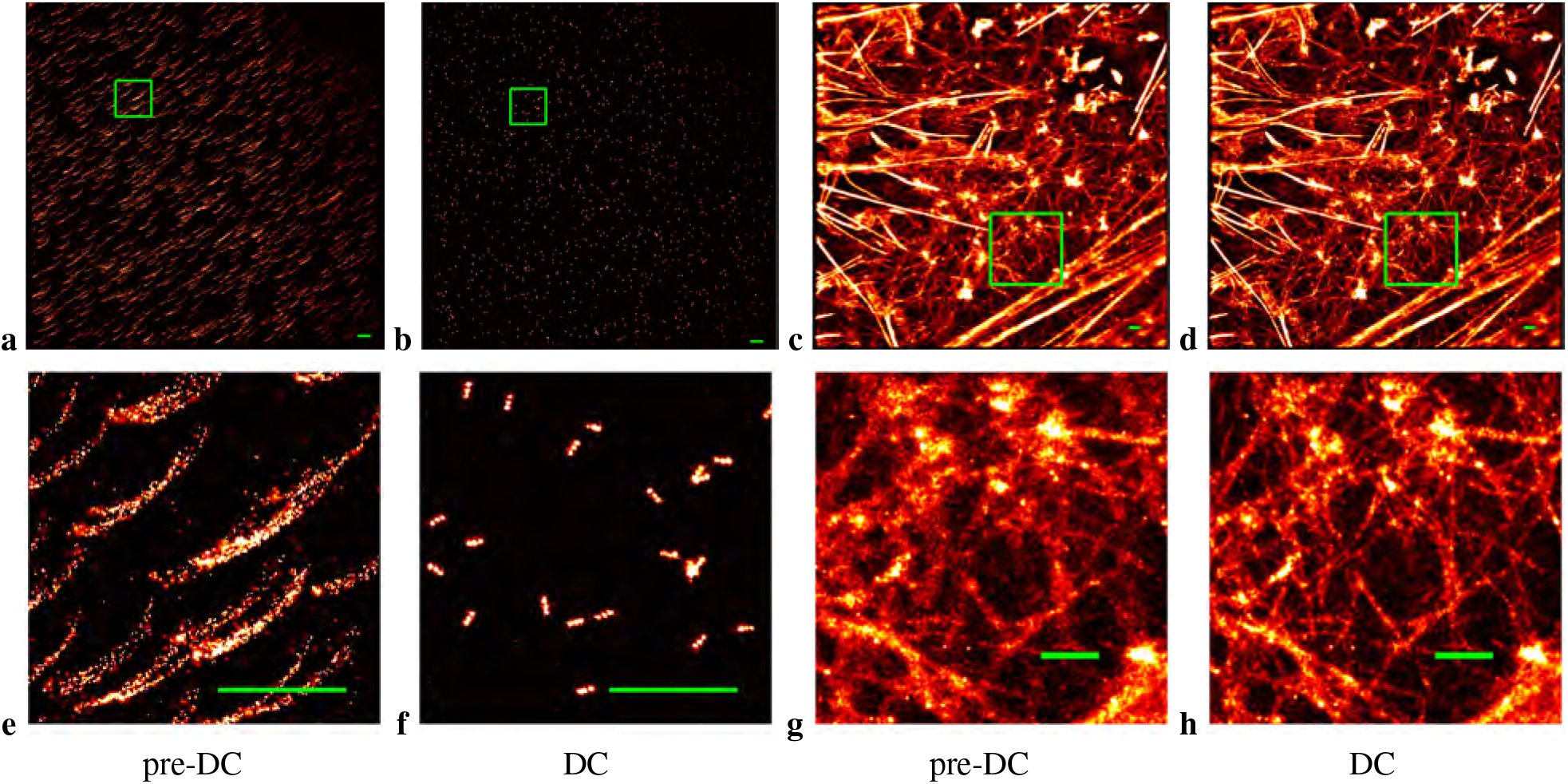
2D drift correction for examples of DNA-PAINT and reversibly binding Lifeact localizations. (a,b) 80 nm nanorods with spots 40 nm apart produced by DNA-PAINT. (e,f) Zoomed in view of the selected region in the previous images. (c,d) Actin microfilaments in HeLa cells. (g,h) Zoomed in view of the selected region in the previous images. (a,c,e,g) Pre-Drift corrected (pre-DC) image. (b,d,f,h) Drift corrected (DC) image. All scale bars measure 1 *μ*m.

The estimated drift plots for the above examples are displayed in Supplementary Fig. S1. The computed drift for each dataset taken for a super-resolution image is plotted with a tapering line segment indicating increasing frame number (so increasing time), while all the frames in the entire image are color coded from blue to red, again indicating the direction of increasing time. The DNA nanorods example did not employ brightfield registration, while the actin example did, therefore, the inter-dataset fitting optimization for each dataset was initialized with either the computed drift correction determined for the last frame in the previous dataset or zero, respectively. These initializations mimicked the drift correction results. The plot for the DNA nanorods example shows a connected series of drift corrections, while the actin example shows drift corrections initially scattered about (0, 0), then slowly increasing in *x* over time.

The improvements in average resolution can be quantified using a measure based on Fourier ring correlation (FRC)^29^. Supplementary Figure S2 shows the results of applying *Q*-corrected FRC (which removes spurious correlations) to the two example datasets where the resolution is defined as the inverse of the spatial frequency when the FRC drops below the black line representing the 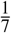 threshold. See **Methods** for more details. Drift correction improved the resolution of the the DNA nanorods example (over the entire ROI) with respect to the uncorrected result from 97.2 nm to 28.7 nm, while for the actin example, the resolution improved from 72.5 nm to 60.9 nm.

In Supplementary Fig. S3, we plot cost function landscapes explored by the optimizer for the two 2D examples presented in Fig. 3. The cost functions for the default fits, linear with zero constant term intra-dataset or constant separation inter-dataset, depend on only one *x* and one *y* parameter, and so can be plotted as 3D surfaces as a function of these parameters. The two 2D examples are displayed in separate rows, exhibiting the initial intra- and inter-dataset landscapes. In all cases, the cost function landscape is funnel shaped with a broad plateau at the top. For the intra-dataset figures, the tip of the funnel is sharp and very near (0, 0), while for the inter-dataset figures, the funneling behavior is more gradual and broad. These landscapes imply that the initial guess should not be too far from the coordinates of the funnel’s bottom, but if it is anywhere close, the funnel will lead the optimizer to the global minimum very quickly.

### Dependence on labeling density and blinking statistics

To better understand the behavior of our drift correction algorithm, we evaluated the performance of estimating artificial drift under varying numbers of particles per dataset. Supplementary Figure S4 shows the results for two studies, one involving randomly generated emitters confined to a 2D star-shaped domain, and the second involving a uniform distribution of random emitters over the entire 2D ROI. The mean drift computed by the algorithm matches within 0.1 (0.05) nm/frame the actual lateral shift imposed upon the emitters in the two scenarios down to about 10^2^ (10^3^) emitters per dataset for star (uniformly) distributed data. The variability of the results, however, broadens dramatically for sparser and sparser datasets around these values. Supplementary Figure S5 provides a set of typical examples for the 2D star shaped domain undergoing artificial drift and then being corrected for a subset of the particle numbers per dataset displayed in Supplementary Fig. S4. The last row in this figure shows the corresponding estimated drift plots for this data as described earlier.

The intra-dataset drift correction procedure depends on nearest neighbor pairings of “on” fluorophores that lie within a distance threshold. We performed a series of simulations for various emitter properties on single 1,000-frame datasets of uniform randomly distributed sets of 2D emitters over a fixed size region in which only intra-dataset drift correction occurs, applying a constant drift per frame. The first two graphs in Supplementary Figure S6 plot the root-mean-square error (RMSE) between the true and the estimated drift curves against the number of blinking event pairs in the dataset, *N_p_N_e_,* where < *N_P_* >= *λ*^2^/2 (see **Methods**) is the expected number of pairs of blinking events per emitter, *λ* is the expected number of blinking events per emitter, and *N_e_* is the number of emitters in the dataset, all calculated from the simulated data. The first simulation series holds *N_e_* fixed with *λ* increasing, while the second holds *λ* fixed with *N_e_* increasing. As the number of pairs of blinking events increases in either situation to 10^2^–10^3^, the RMSE drops precipitously to about 1 nm. See also **Supplemental Methods; RMSE analysis** for additional details.

Plotting the theoretical product (see **Methods; Fluorophore pairings for intra-dataset drift correction**) of < *N_p_* > *N_e_* over a range of *λ* and *N_e_* produces a three-dimensional surface (Supplementary Fig. S6), which can also be displayed as a contour plot with contours of constant < *N_p_* > *N_e_*. This coupling of the plots in Supplementary Fig. S6 constrain the values of *λ* and *N_e_* for producing a desired RMSE under this particular intra-dataset model.

In Supplementary Fig. S6, the simulations considered a linear model for the drift and applied a linear polynomial fit for the correction estimation procedure. In Supplementary Fig. S7, the results for other studies are shown. A linear drift was approximated, using about 10^3^-10^4^ pairs for 1 nm accuracy, by a quadratic fit (but was about 5× slower than the case for a linear fit), a quadratic drift was approximated by a quadratic fit also using approximately 10^3^–10^4^ pairs for nanometer accuracy, and a quadratic drift was less well approximated by a linear fit, achieving asymptotic 3–5 nanometer accuracy starting at 10^2^-10^3^ pairs.

### 3D simulations

A more complex example is provided in Fig. 4. Here, two 3D rings of diameter 40 nm containing localizations are separated by 80 nm. A simulated astigmatism point spread function (PSF) was used to generate the localizations. Smooth, slowly oscillating drifts are applied in the *x, y* and *z* directions for 50,000 frames, maximally varying approximately 200 nm over about 12,500 frames. The simulated data is broken up into 100 datasets of 500 frames apiece. Applying the post-processing drift correction algorithm to the drifted data reproduces the rings with some slight scatter in the *z*-direction (see Fig. 4(a,b,c)). The computed drift curves almost perfectly overlap the simulated drift. Figure 4(j,k) show, respectively, the differences between the computed and simulated drift curves, and the overlapped *y*-drift curve. The greatest differences tend to occur around steep changes in the derivative of the drift curve. A second study in which 10 ring pairs were centered at different *x, y, z*-locations was performed using the same drift curves. The results are nearly identical to those for the single ring pair (see Fig. 4(d,e,f,l)), even for a zoomed-in set (Fig. 4(g,i)). The estimated drift plot, Fig. 4(h), is comparable in shape to the drifted dataset from the first study, Fig. 4(b).

**Figure 4.**
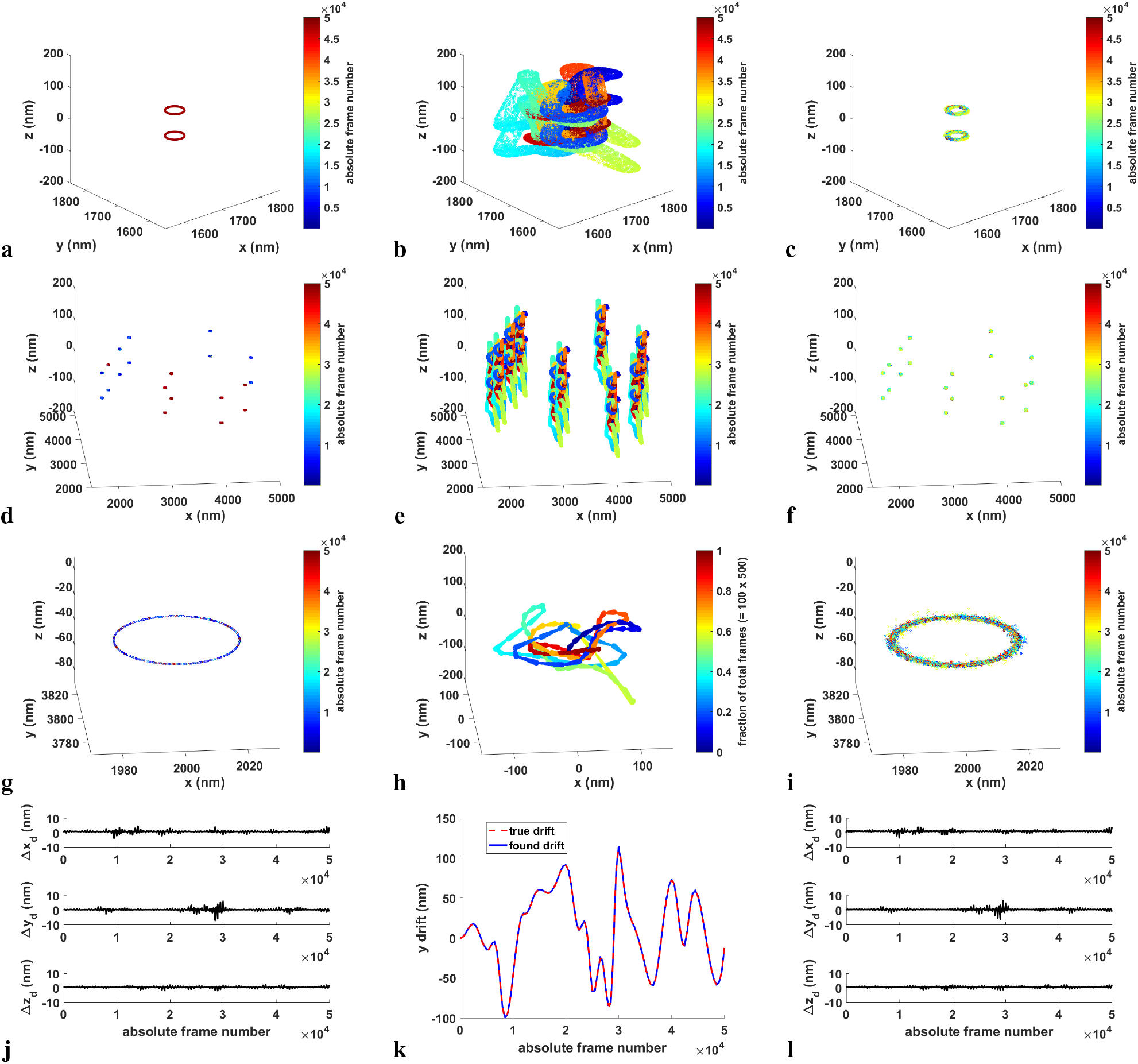
3D drift correction applied to simulated 40 nm diameter rings separated by 80 nm. Simulated PSFs and drift curves were used. 50,000 frames were generated, divided into 100 datasets of 500 frames each. (a) True image of the original random emitters confined to the two rings. (b) Drifted image. The emitters are color-coded by frame number. (c) The drift-corrected image produced by driftCorrectKNN using default settings. (d,e,f) True, drifted and drift-corrected images in which multiple (10) pairs of rings centered at different *x, y, z*-positions were simulated. (g,i) Zoomed-in true and drift-corrected ring pair (lowest one shown in (d,f)). (h) Corresponding estimated drift plot for the multiple ring pair example. (j,l) The differences between the estimated and simulated *x, y, z*-drifts as a function of the absolute frame number for (j) the single pair of rings example, (l) the multiple ring pair example. (k) Estimated versus simulated *y*-drift for the single pair of rings example.

The results with the above 3D localizations were quite good. However, we decided to perform a pair of studies with noisier simulated data, in which the localizations were fit and thresholded (see **Methods**) after applying drift to them in order to mimic more realistic results. The first set of localizations were generated as a pair of 80 nm separated rings using a simulated astigmatism PSF and the same drift curves as before. The results are shown in Supplementary Fig. S8(a,b,c,g,h). The data is much more scattered, but ring separation is still clear. However, if we used an experimental PSF for simulation and fitting (**Methods**), 80 nm separation was not sufficient to distinguish the rings, so the results shown in Supplementary Fig. S8(d,e,f,i) are for a pair of rings separated by 120 nm. The differences in computed versus simulated drift curves are also larger for the imperfect data, especially when using the experimental PSF (Supplementary Fig. S8(g,i)).

### Breaking up datasets

Finally, sometimes it is beneficial to reorganize the acquired datasets when no brightfield registration has occurred. Supplementary Figure S9 shows an example of ATTO655 20 nm nanorulers in which the post-processing drift correction procedure was applied with varying dataset organizations. Here, 15,000 frames in a single dataset provided only one opportunity for the intra-dataset and no opportunities for the inter-dataset portions of the drift correction algorithm to be applied, so the results were very smeared out like the raw super-resolution image, while 100 frames per dataset provided too few localizations per dataset for clean sum images to be used by the algorithm. 1,000 frames per dataset produced the sharpest results. We find 500-1,000 frames per dataset to be a good choice for typical drift correction on single molecule localization microscopy (SMLM) data.

## Discussion

To correct super-resolution drift, we propose a combination of brightfield registration performed after each dataset acquired, and a post-processing algorithm based on nearest neighbor distances, combining intra-dataset and inter-dataset processing. No extra hardware is required. This methodology can produce a significant improvement in sharpness as indicated by FRC results for real data, while theoretical results and simulations show under what fluorophore conditions nanometer accuracy can be achieved. The procedure works in both 2D and 3D, the brightfield registration severely curtailing the inter-dataset drift and the post-processing algorithm then correcting for whatever drift remains.

Our post-processing drift-correction algorithm works for images produced by common super-resolution methods such as dSTORM and DNA-PAINT in which fluorescent labels blink or dye-labeled DNA strands bind/unbind repeatedly. In this situation, the nearest neighbors provide a cheap and fast way to produce image feature pairings that remain roughly stable over a number of frames. Note that a continuous underlying structure is not required, as the algorithm also works for isolated and randomly distributed fluorophores. However, this idea will not work with single activated probes such as in photo-activated localization microscopy (PALM)^11^. Minimizing the thresholded sum of the nearest neighbor distances keeps the localizations, and thus the emitters, cohesively moving together over time. A bad model would let these distances spread out and the emitters drift apart contrary to the assumptions of the model.

The intra-dataset algorithm estimates drift as a function of frame number, while the inter-dataset algorithm computes lateral shifts between datasets. Increasing the number of blinking events in the intra-dataset algorithm provides more true pairs to constrain the drift model.

The intra-dataset threshold on nearest neighbor distances minimizes the effect of false fluorophore pairing. A threshold is less important for inter-dataset fitting as the sum of all the emitters in entire datasets rather than the sparse representation present in individual frames are considered when computing nearest neighbor distances, however, it can be important for sparse datasets. Employing thresholds is in contrast to cross-correlation techniques where contributions from different emitters are not restricted, so that image overlap at a pixel may be due to a combination of near and far away emitters. The nearest neighbor thresholds restrict contributions to local entities. Note that Eq. 1 is reminiscent of the MCF function^5^, but with a simpler structure and thresholding present, so significantly faster computationally. The threshold paradigm allows up to nanometer accuracy (for the intra-dataset portion of the algorithm) under practical conditions.

## Methods

### Sample preparation

HeLa cells (ATCC) were cultured as described in detail previously^30^. All labeling and washing steps were carried out at room temperature. Cells were seeded onto 25 mm glass (#1.5) coverslips in 6 well chambers (LabTek) to adhere for 24 h. For alpha tubulin, we used PBS as a buffer, while a PEM buffer (80 mM PIPES + 5mM EGTA + 2mM MgCl2, at pH 7.2) was used for actin filaments. The first fixation step was in 0.6% paraformaldehyde + 0.1% glutaraldehyde + 0.25% Triton diluted in buffer for 60 seconds, followed by a hard fixative buffer including 4% paraformaldehyde + 0.2% glutaraldehyde diluted in buffer for an hour. Cells were washed 2x in PBS and kept in NaBH_4_ for 10 min to quench the autofluorescence within the cells resulting from glutaraldehyde in the fixation buffer, followed by a 2x wash with PBS. To quench reactive crosslinkers, the samples were kept in 10 mM Tris for 10 min, followed by 2 washes with PBS. Finally, samples were blocked in 5% BSA + 0.05% Triton X-100 for 15 min. At the end, samples were washed 1x with PBS.

For alpha tubulin, fixed cells were immunolabeled using Alexa Fluor 647 conjugated primary antibody (Novus Biologicals, CO) at 10 *μ*g/mL for 1hr, followed by washes using PBS. For actin filaments in HeLa cells, the sample was imaged while it was labeled by Lifeact + Atto 655 with 0.1% BSA to minimize non-specific intracellular binding of the Lifeact + Atto 655. The labeling and imaging buffer was 3 nM of Lifeact + Atto 655 in 50 mM Tris, 10 mM NaCl, 10% (w/v) Glucose (TNG) at pH 8. GATTA-PAINT nanoruler slide sample (40R, GATTAquant DNA Technologies) was used as purchased.

### Imaging

For alpha tubulin in HeLa cells and the DNA-PAINT nanorods sample, imaging was done on a custom-built inverted wide field fluorescence microscope setup as described previously^30^. Fluorescence excitation of the sample was done using 642 nm laser diode (Thorlabs, HL6366DG). The laser beam was collimated and passed through a single mode fiber (Thorlabs, P1-488PM-FC-2) before being focused on the back focal plane of a 1.45 NA oil objective (UAPON 150XOTIRF, Olympus America Inc.). TIRF excitation of the sample was achieved by translating the laser close to the edge of the objective back aperture. Fluorescence emission collected from the sample was passed through a quad band dichroic/emission filter set (Semrock, LF405/488/561/635-A) and a bandpass filter (Semrock, FF01-446/523/600/677-25) before being detected using an electron-multiplying charge-coupled device (EM CCD) camera (Andor Technologies, iXon 897). Data collection and instrument control on the microscopes here and below was controlled by custom-written MATLAB software^31^.

For dSTORM imaging of alpha tubulin in HeLa cells, an xyz piezo stage (Mad City Labs, Nano-LPS100) mounted on an x-y manual stage was installed on the microscope for cell locating and brightfield registration. To mount the prepared samples on 25 mm coverslips, an Attofluor cell chamber (Life Technologies, A-7816) was used and a clean 25 mm coverslip was used to seal the samples. A trans-illumination halogen lamp equipped with the microscope was used for collecting the brightfield images. The samples were stored at 4°C and then transferred to the microscope just before starting imaging at room temperature. This natural heating up of the sample to room temperature was used to create the sample drift, which was observed in the images. A total of 50,000 256 × 256 pixel frames were collected using 100 ms exposure time for the GATTA-PAINT nanoruler super-resolution imaging. For alpha tubulin in HeLa cells, 25,000 frames (256 × 256 pixels) were recorded in each imaging experiment using 10 ms camera exposure time.

For actin filaments in HeLa cells, the imaging system was built on an inverted microscope (Olympus). An xyz piezo stage (Mad City Labs, Nano-LPS100) mounted on a x-y manual stage was installed on the microscope for cell locating and brightfield registration. A mounted LED with wavelength 850 nm (M850L3, Thorlabs) was used for brightfield illumination. Brightfield images were collected on a complementary metal-oxide semiconductor (CMOS) camera (Thorlabs, DCC1545M) after reflection by a short-pass dichroic beam splitter (Semrock, FF750-SDi02) and passing through a single-band bandpass filter (Semrock, FF01-835/70-25). A 638 nm laser was used (collimated from a laser diode, Thorlabs, L638P200) coupled into a single mode fiber and focused onto the back focal plane of the 1.49 NA objective lens (Olympus UAPON 100XOTIRF). Emission for super-resolution data was collected through a short-pass dichroic beam splitter (Semrock, FF750-SDi02) and a single-band bandpass filter (Semrock, FF624-Di01) on an iXon 860 EM CCD camera (Andor Technologies, iXon DU-860E-CS0#BV).

Imaging was performed in TNG buffer mixed with Lifeact conjugated with dye. The blinking events for single molecule localization were due to the binding/unbinding of Lifeact to the actin filaments^32,33^. We used the same 25 mm coverslip chamber as above to mount the samples on the microscope. Imaging was performed with TIRF illumination to reduce the background noise due to diffusing dyes in the imaging buffer, and images were acquired at 20 ms exposure time. Brightfield registration was performed to correct for drift after every 3,000 frames as mentioned in **Methods; Brightfield registration**.

### Super-resolution fitting

We used a difference of Gaussians filter to reduce noise and enhance the signal from single emitters to enable identification of local maxima. Pixel coordinates with local intensity maxima were selected and used as centers of fitting regions of size 16 × 16 pixels. A given numerical PSF was used in the GPU 3D fitting algorithm to localize emitters. The algorithm fit the single emitters using maximum likelihood estimation (MLE) and assuming a Poisson noise model^34,35^. The algorithm finds the MLE employing the Newton-Raphson approach to iteratively update parameters, including *x, y, z*-locations, intensity and background. The resulting localizations were filtered by thresholding the intensity, background, *p*-value and Cramer-Rao lower bound of the estimations.

### Brightfield registration

A camera, a stage, and a lamp are the hardware required to perform brightfield registration. At the beginning of the experiment, a brightfield reference *z*-stack is produced after turning the lamp on (the lamp is normally turned off during the collection of movie frames to not interfere with the fluorescent particle emissions). A dataset is then collected, after which a brightfield image *z*-stack is produced to be compared with the reference *z*-stack, again turning the lamp on during its production. The *z*-stack in our setup consists of 2D images taken at an odd number (typically 21) of different *z* positions reachable by the stage. Aligning the current *z*-stack with the reference *z*-stack, an (*x, y, z*) drift is computed for it, which will then be corrected before the next super-resolution dataset is imaged. The process then repeats until the found shifts in *x, y* and *z* are all less than a selected tolerance (typically chosen to be between 5–50 nm, depending on the minimum step size of the stage). The brightfield registration procedure typically takes < 30 seconds per super-resolution dataset collected.

The software used to perform brightfield registration is included in the Supplementary Software.

### Post-processing drift correction (driftCorrectKNN)

driftCorrectKNN operates by taking (*x, y*) or (*x, y, z*) coordinates from a sequential series of image frames of super-resolution observations collected together in datasets and computes the drift correction over the frames.

The intra-dataset drift correction is computed by optimizing a polynomial fit as a function of frame number (so really time) using Eq. 1 with *l* replaced by *l_intra_* = 1 pixel. In practice, a linear polynomial fit was sufficient to get an accuracy of 0.0001 pixel/frame (~0.01 nm/frame) for the drift rate. Finding more than one nearest neighbor did not noticeably improve the results.

The inter-dataset drift correction is computed by optimizing a constant fit to the positions of all localizations in each dataset relative to corresponding drift-corrected positions computed for the initial dataset in the intra-dataset analysis. Eq. 1 with *l* replaced by *l_inter_* = 2 pixels applies once again in computing the sum. This is done rather than considering the nearest neighbor distances between the predicted positions in the current dataset (derived from the initial dataset and the modeled drifts) and the actual positions. The two points of view are very similar, however, the former requires only one *k-d* decomposition (of the initial dataset), while the latter viewpoint requires a *k-d* decomposition of every dataset but the first, so is much more computationally demanding. If no other phenomena but drift is happening and the drift model is accurate, this nearest neighbor sum should be zero.

The values chosen for *l_intra_* (1 pixel) and *l_inter_* (2 pixels) were the result of a parameter study involving drift correction on the images and simulations presented in this paper. We found it necessary to make *l_inter_* > *l_intra_*, and the values chosen seemed to give reasonable overall performance.

The choice of using zero or the ending drift corrected value of the previous dataset, as derived from the intra-dataset and previous inter-dataset drift corrections, to initialize the inter-dataset optimization depends on how the data was collected. If brightfield registration was performed, then zero (correct for the first dataset, and the sum of accumulated drifts, so theoretically zero, in subsequent datasets) is appropriate. If the datasets were not collected using registration, then the ending frame of the previous dataset adjusted for accumulated drift is the better choice for initializing the inter-dataset optimization. The intra/inter-dataset optimizations, which use the thresholded/simple nearest neighbor sums above as cost functions, search for the global minimum of these costs using the MATLAB function fminsearch.

If each dataset in an experiment contains a large number of frames in which drift changes are significant and the datasets have not been registered, breaking up the datasets into smaller chunks can sometimes help to produce better results. The user can specify the total number of datasets or number of frames per dataset desired while calling driftCorrectKNN and the code will internally reorganize the datasets as specified, perform the drift correction calculation, and then reassemble the results back into the original dataset scheme at the end (see, for example, Supplementary Figure S9).

driftCorrectKNN is implemented in MATLAB (see **Supplemental Methods; Post-processing drift correction algorithm** for a MATLAB pseudocode description) and is included in the Supplementary Software.

### Fluorophore pairings for intra-dataset drift correction

The intra-dataset drift correction algorithm computes the sum of nearest neighbor distances (within a threshold) produced by the pairings of nearby “on” fluorophores over an entire dataset. The sum of these distances is the cost function, which when minimized, produces an estimate of the drift as a function of the frame number. To understand this pairing phenomenon in more detail, consider for a dataset how one might compute the expected number of pairs of blinking events per emitter, < *N^p^* >, given the number of expected blinking events per emitter, *λ*, which, for example, can be directly computed in simulations. We examined simulations exhibiting a uniform distribution (see **Methods; Simulations**), where single datasets of 1,000 frames were used, so that only intra-dataset drift correction was performed. The number of blinking events post-frame connection was computed for the various emitter densities averaged over N = 100 runs.

Let *P(k; *λ*)* be the probability of exactly *k* blinking events (*k* ≥ 0) per emitter occurring given an expected value of *λ* blinking events per emitter. This quantity can be represented by the Poisson distribution *P(k;λ) = λ^k^e^-λ^/k!*. The number of blinking event pairs per emitter, *N_p_(k)*, given *k* blinking events, is simply the number of ways of choosing two blinking events from a total of *k* available, 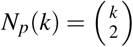. Therefore, the total number of pairs of blinking events per emitter expected in the dataset is

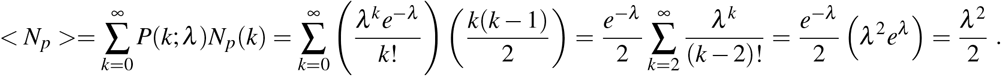

### Fourier ring correlation (FRC)

FRC^29^ is a measure of the average resolution over a super-resolution image. It works by dividing the set of single-emitter localizations in the super-resolution image into two statistically independent subsets. The Fourier transforms of subimages generated from each of these subsets are then statistically correlated over pixels on the perimeters of circles of constant spatial frequency. The image resolution is defined as the inverse of the spatial frequency when the FRC curve drops below a threshold, taken to be 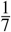 in Nieuwenhuizen et al.^29^. See Supplementary Fig. S2. Spurious correlations (for example, due to repeated photoactivation of the same emitter) are removed by estimating the number of times an emitter is localized on average (*Q*) assuming Poisson statistics. All analyses were accomplished using the software developed by Nieuwenhuizen et al. found at http://www.diplib.org/add-ons.

### Simulations

2D simulations of simple artificial drift were performed for (noisy) emitters randomly distributed in a star-shaped domain and over the entire 6400 × 6400 nm^2^ ROI. Localization uncertainties around the true emitter locations varied by up to ± 10 nm. Seven different initial fluorophore densities were specified ([10, 5, 2, 1, 0.5, 0.25, 0.125] · 10^-4^fluorophore/nm^2^) from which the number of emitters per dataset were computed. A constant drift of Δ*x* = 0.3 and Δ*y* = 0.4 nm/frame was imposed upon each emitter, after which driftCorrectKNN was applied to the drifted results. *N* = 100 simulations per condition were performed and the mean resulting drift calculated was compared to the true drift. The standard deviations of the results were also computed.

To produce the RMSE vs. fluorophore pairings plot (Supplementary Fig. S6(a)), we performed simulations in which emitters were randomly distributed uniformly over a 25,600 × 25,600 nm^2^ ROI using single 1,000 frame datasets, allowing for only intra-dataset drift correction. The following two series were run for linear and quadratic drifts. The first varied *λ*, the number (via a Poisson distribution) of expected blinking events per emitter per dataset over the range 0.01-10 while fixing the number of emitters, *N_e_*, to 10^5^. The second series varied *N_e_* over 10^3^-10^6^ while setting *λ* = 0.2. True cumulative drifts were defined by *αf* + *βf*^2^, where *f* is the frame number. Linear drifts were *α*_*x*_ = 0.05 nm/frame and = 0.02 nm/frame, while quadratic drifts had the above linear terms as well as quadratic terms *β*_*x*_ = 10 nm/frame^2^, *β*_*y*_ = −20 nm/frame^2^. driftCorrectKNN was run with the appropriate intra-dataset polynomial order. The results were averaged over *N* = 100 runs.

A second series of simulations involved constructing more realistic artificial drift curves, and in general mimicking real, 3D experimental conditions. Simulated drift curves for *x, y* and *z*, designed to resemble realistic drift curves, were produced by generating a set of approximately 35 nodes over 50,000 frames (the exact number of nodes selected from a Poisson distribution), each node representing a point on a graph of drift component value (in units of nm) versus frame number. The intervals between the nodes randomly varied in width with respect to an average of *N*_frames_/(*N*_nodes_ + 1). The drift curves were fit by passing a cubic spline through the nodes in which the component values ranged approximately between −100 to +100 nm (−1 to +1 pixel). See Fig. 4(k) for an example of one such drift curve, the *y*-component.

In the first simulations, two 40 nm diameter rings separated in the *z*-direction by 80 nm were produced to mimic a DNA-PAINT labeled nanorod in which the center and the two ends were visible under super-resolution. The 2D sample structure, a ring in this case, was given as a binary image template, where the nonzero portion was filled with emitters of a uniform density *ρ* = 1,000 fluorophores/pixel unit of length. A trace of blinking events for each emitter was then generated using the input duty cycle parameters, *K*_on_ = 0.0005/frame and *K*_off_ = 1/frame, which are the rate of emitters turning from off to on and from on to off. The random durations for the emitters to turn on and the on-durations were taken from exponential distributions with mean values of, respectively, *K*_on_ and *K*_off_. The density of the on-emitters was then given by

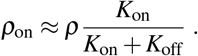

The simulated structures were produced at specific *z*-positions, which allowed separated pairs of rings to be made. A second set of simulations involved 10 pairs of 80 nm separated rings randomly distributed in (*x, y*) around the simulation region. A theoretical astigmatism point spread function (PSF) was used for the 80 nm separated sets, while a spiral experimental PSF was given as input for a third set of simulations in which a single pair of rings separated by 120 nm was studied. Localizations in the third set of simulations were further fit and thresholded (see **Methods; Super-resolution fitting** for additional details). For all simulations, an interpolated PSF was generated according to the *z*-position of the emitter and placed in (*x, y, t*) in the absence of drift. Drift was added as discussed above. If an emitter was on for the entire exposure time of a frame, it was taken to be at maximum intensity (12,000 photons), otherwise dimmer based on the fraction of the frame in which it was on. The data was produced with a fixed offset background (1,000 photons) corrupted with Poisson noise.

## Data availability

The DNA-PAINT 80 nm nanorods example is included in the Supplementary Software. All datasets analyzed during the current study are available from the corresponding author on reasonable request.

## Acknowledgments

This work was supported by NIH grants NIBIB 1R21EB019589 and NIGMS 1R21GM132716, and the New Mexico Spatiotemporal Modeling Center (NIH P50GM085273). Some of the development and computations were performed at the Center for Advanced Research Computing, supported in part by the National Science Foundation, at the University of New Mexico. Farzin Farzam developed a previous drift correction code.

## Author Contributions

HMF and SP provided experimental data. DJS and HMF developed the 3D brightfield registration algorithm and the associated microscope control software, respectively. HMF implemented the frame connection algorithm. MJW and KAL developed the post-processing drift correction algorithm. MF developed the GPU 3D fitting codes. MJW and MF performed the simulations. MJW wrote the manuscript with contributions from the other authors. All authors reviewed the manuscript.

## Additional Information

**Supplementary information** accompanies this paper.

**Competing Interests:** None of the authors declare competing interests.

---

**Figure S1.**
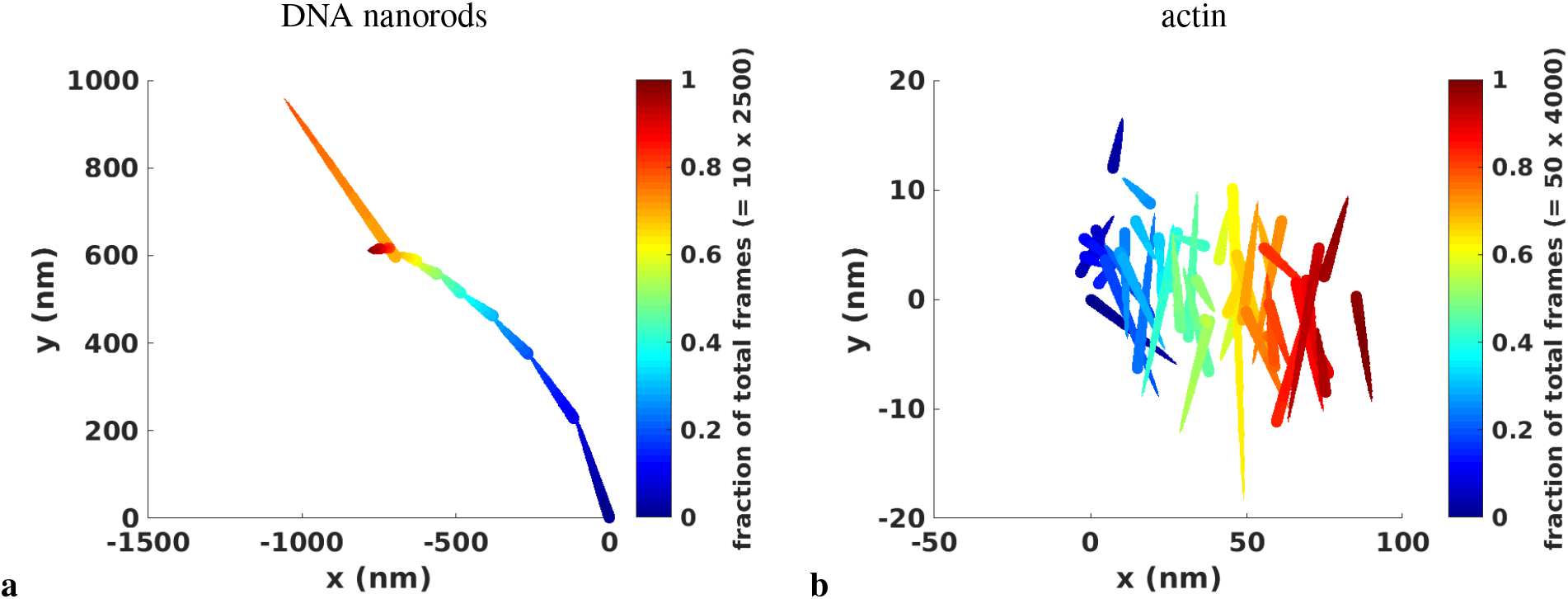
2D estimated drift plots. Each dataset is represented by a separate line segment. Frames are color coded from blue to red to indicate the passage of time. Within each dataset, passage of time is also indicated by the segment width tapering from large to small (like an arrowhead). (a) 80 nm nanorods with spots 40 nm apart produced by DNA-PAINT. The initial guess used for the inter-dataset optimization procedure was the drift-corrected values from the last frame (2500) of each previous dataset. (b) Actin microfilaments in HeLa cells. The initial guess used for the inter-dataset optimization procedure was zero.

**Figure S2.**
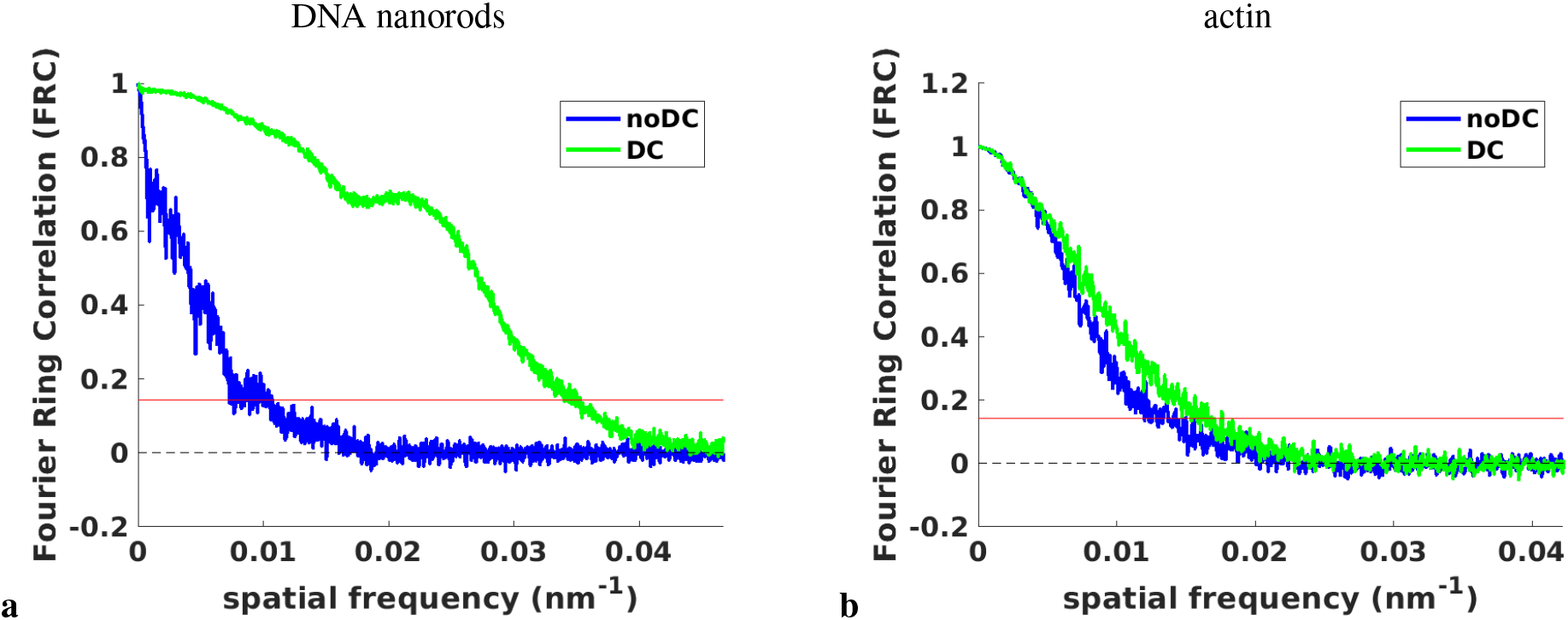
Fourier Ring Correlation plots. (a) 80 nm nanorods with spots 40 nm apart produced by DNA-PAINT. (b) Actin microfilaments in HeLa cells. The blue lines represent raw (uncorrected) data, while the green lines are the results given data drift corrected through the post-processing algorithm. The black horizontal line at 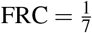 is the threshold that when crossed by the FRC curve defines the spatial resolution of the image, as the inverse of the spatial frequency at the crossing point. For (a), the estimated resolutions of the uncorrected data compared to the corrected results were 97.2 ± 0.9 nm versus 28.7 ± 0.1 nm, while for (b), these numbers were 72.5 ± 1.4 nm and 60.9 ± 0.6 nm, respectively.

**Figure S3.**
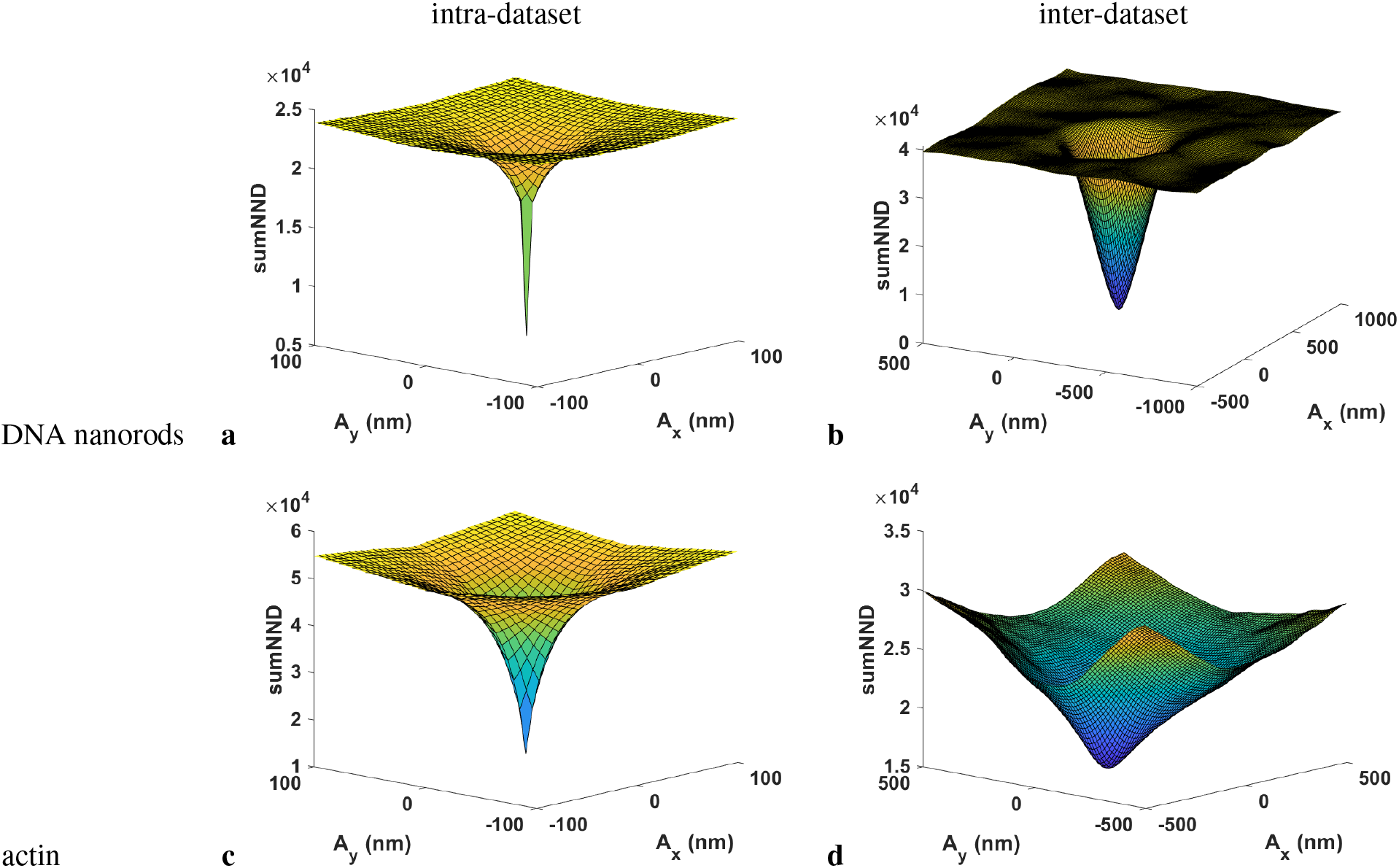
Cost function landscapes for driftCorrectKNN when fitting a linear polynomial with zero constant term to the intra-dataset drift correction and a constant lateral shift to the inter-dataset drift correction. (a,b) Dataset of 80 nm nanorods with spots 40 nm apart produced by DNA-PAINT. (c,d) Dataset of actin microfilaments in HeLa cells. (a,c) Intra-dataset landscape for the first dataset. The (*x, y*) coordinates refer to the coefficients of the linear (and only) term in the polynomial fit of the intra-dataset drift correction. sumNND (defined by Eq. 1) is the cost. (b,d) Inter-dataset landscape for the first dataset processed (dataset 2 which is shifted relative to dataset 1). The (*x,y*) coordinates refer to the constant lateral *x/y* shifts of the inter-dataset drift correction.

**Figure S4.**
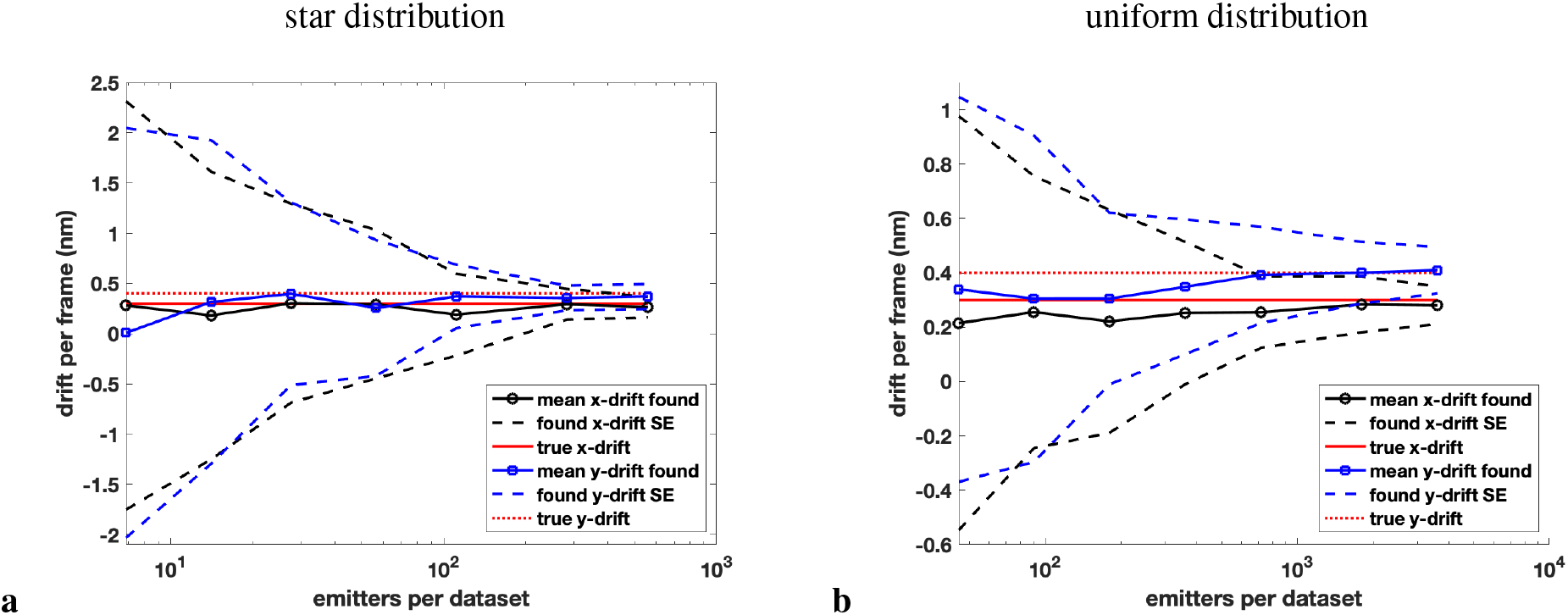
Results of drift correction for true constant shifts applied to noisy random emitters occupying two differently shaped domains as a function of the number of emitters per dataset. (a) 2D star-shaped domain. (b) 2D uniformly distributed domain. 10 datasets of 100 frames each were generated per simulation. A drift of Δ*x* = 0.3 and Δ*y* = 0.4 nm/frame was then applied. The solid black and blue lines are simulation means, while the dashed black and blue lines define one standard deviation about the means. The results were averaged over *N* = 100 simulations for each condition.

**Figure S5.**
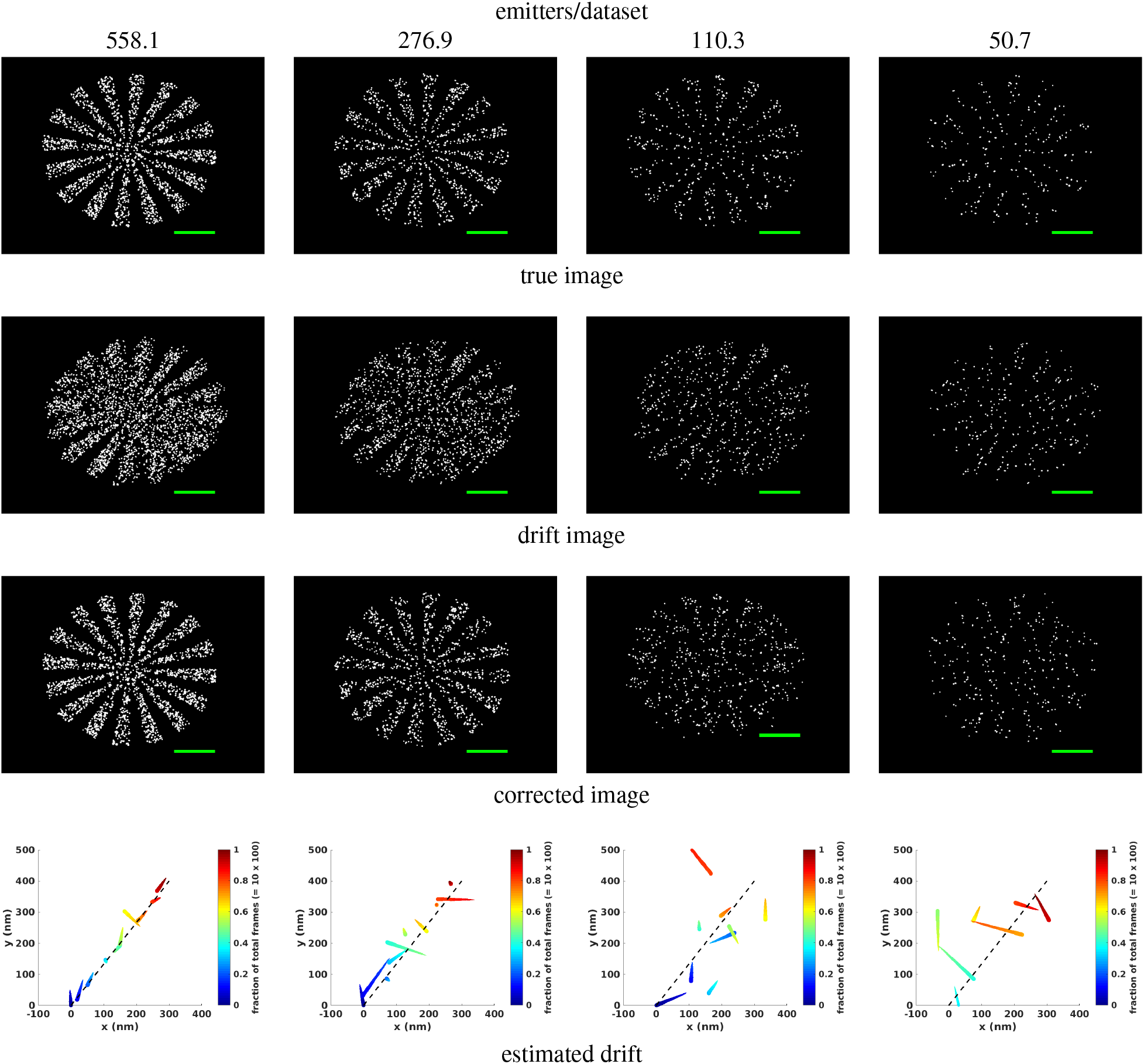
Actions of driftCorrectKNN on a drifted 2D star-shaped domain for various densities of (noisy) emitters per dataset. (1st row) True images of the original random emitters confined to a star-shaped domain. 10 datasets of 100 frames each were generated. (2nd row) Drifted images where a drift of Δ*x* = 0.3 and Δ*y* = 0.4 nm/frame was applied. (3rd row) The drift-corrected images produced by driftCorrectKNN using default settings. (4th row) Estimated drift plots. The black dashed lines depict the true drifts. All scale bars are 1 *μ*m.

**Figure S6.**
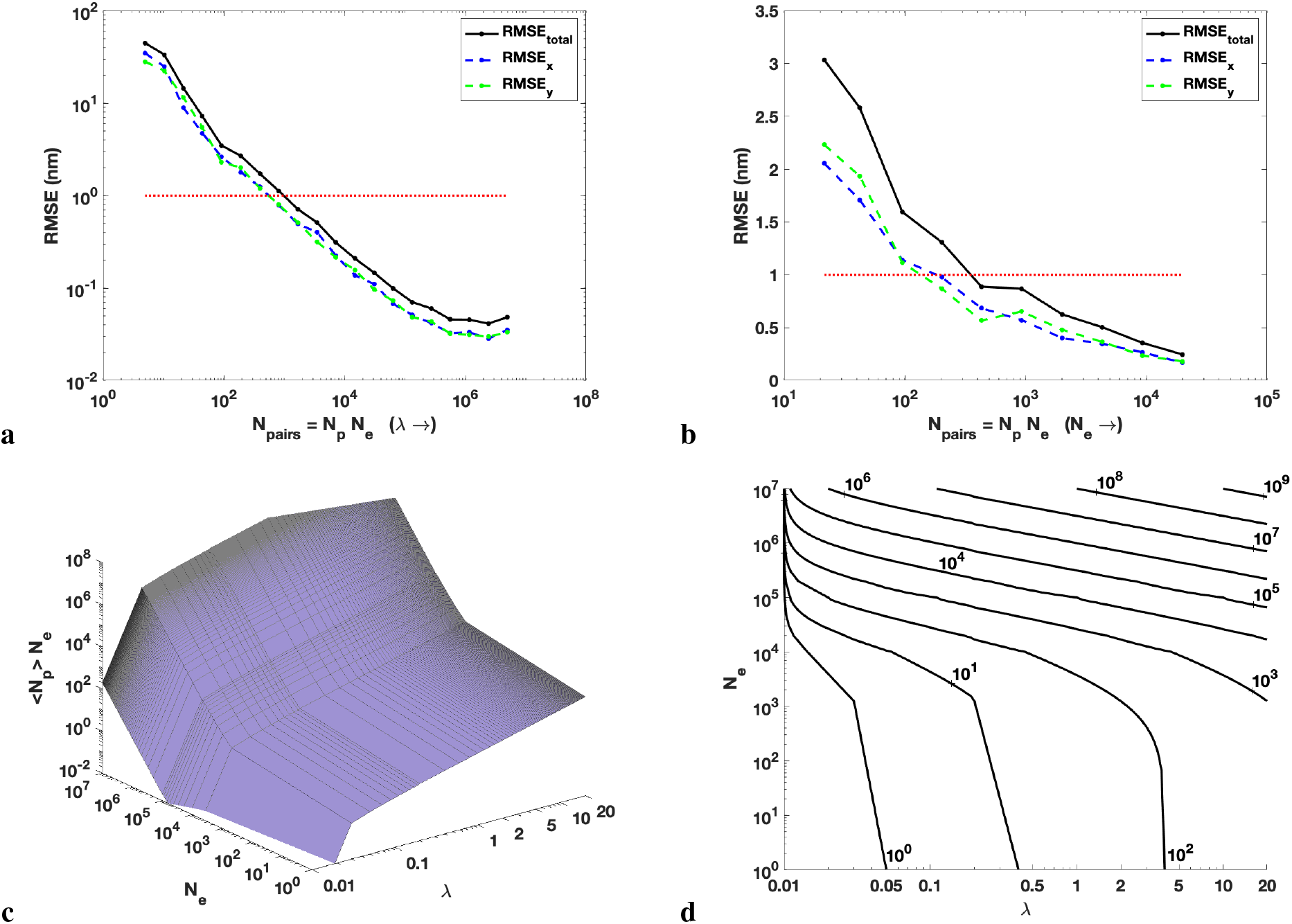
Fluorophore pairings for intra-dataset drift correction. (a,b) RMSE (and *x* and *y* components) between the true and the estimated drift curves plotted versus the number of pairs of blinking events, *N_p_N_e_*, for noisy 2D uniform randomly distributed emitters with (a) *λ* increasing from 0.01–10 and fixed *N_e_* = 10^5^, (b) *N_e_* increasing from 10^3^–10^6^ and fixed *λ* = 0.2. The imposed drift was linear and the intra-dataset fitting was also linear. Results were averaged over N = 100 simulations. The red dotted lines correspond to RMSE = 1 nm. (c) Theoretical 3D plot of < *N_p_ > N_e_* versus the number of expected blinking events per emitter, *λ*, and the number of emitters, *N_e_*, for a dataset. (d) 2D contour plot of the surface in which lines of constant < *N_p_ > N_e_* are displayed.

**Figure S7.**
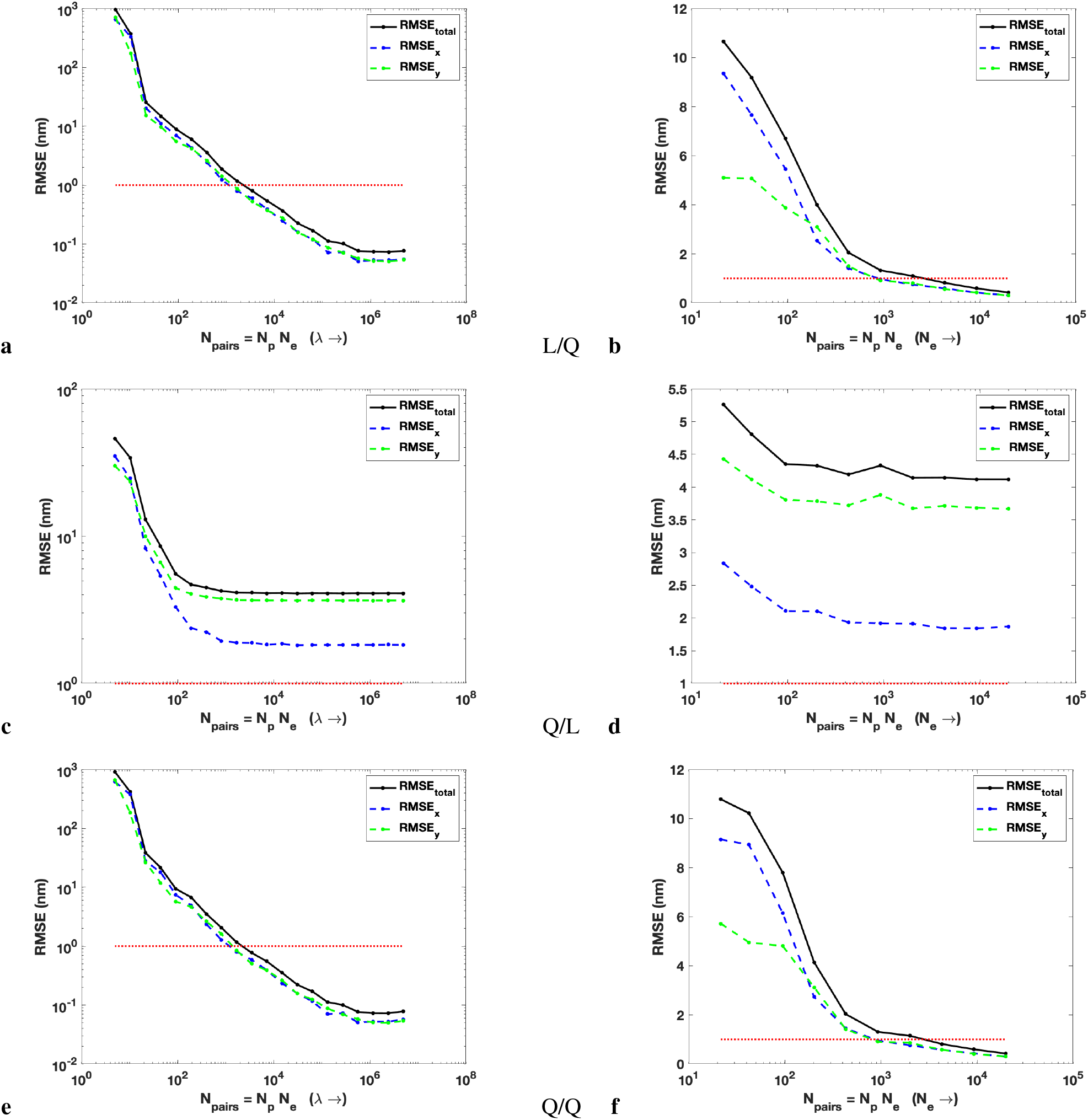
Fluorophore pairings for intra-dataset drift correction. RMSE (and *x* and *y* components) between the true and the estimated drift curves plotted versus the number of pairs of blinking events, *N_p_N_e_*, for noisy 2D uniform randomly distributed emitters with (a,c,e) *λ* increasing from 0.01–10 and fixed *N_e_* = 10^5^, (b,d,f) *N_e_* increasing from 10^3^–10^6^ and fixed *λ* = 0.2. The imposed drift was (a,b) linear / (c,d,e,f) quadratic, and the intra-dataset fitting was (c,d) linear / (a,b,e,f) quadratic. This is also indicated by L/Q (linear imposed drift/quadratic intra-dataset fitting), etc. above. Results were averaged over *N* = 100 simulations. The red dotted lines correspond to RMSE = 1 nm.

**Figure S8.**
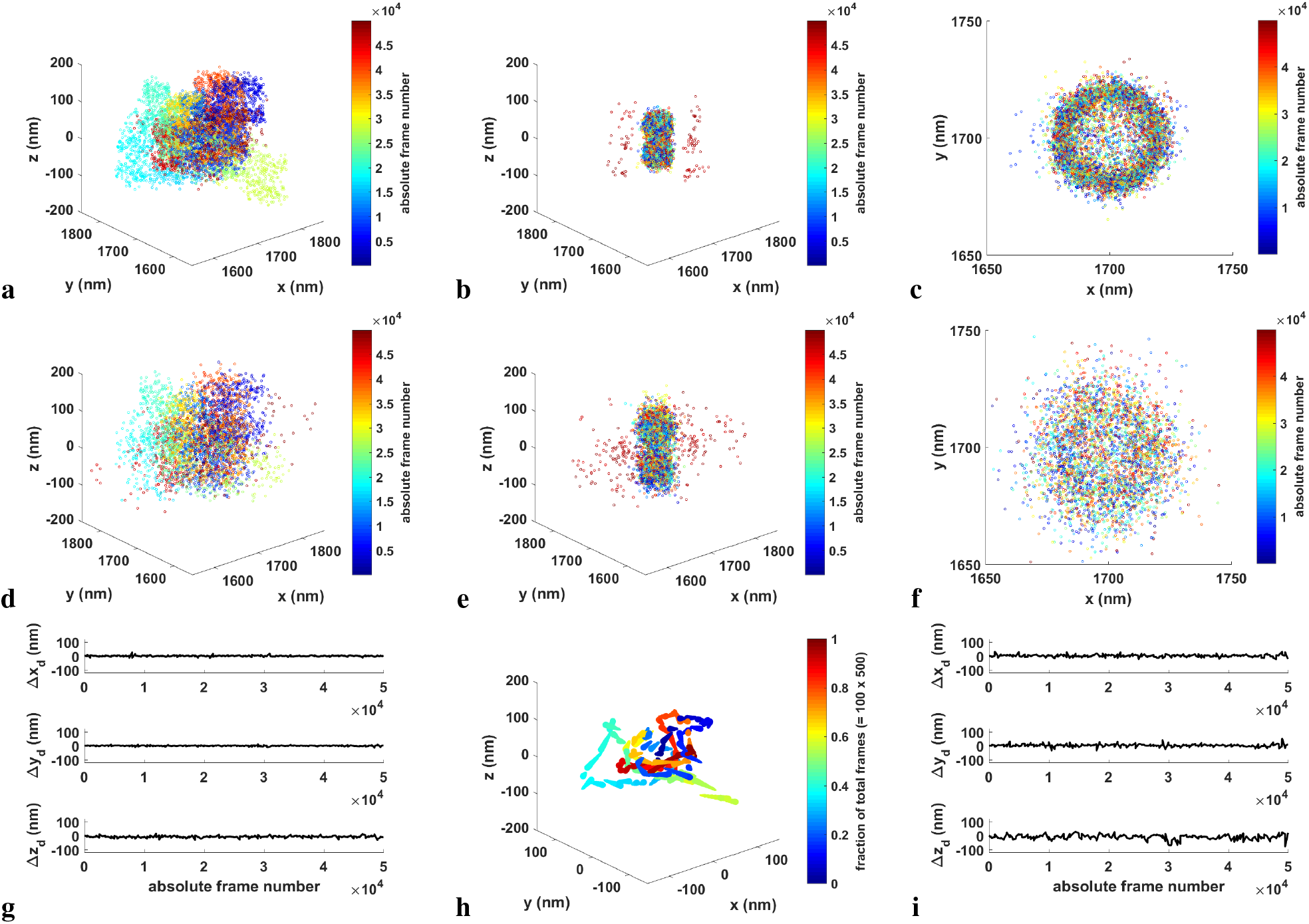
3D drift correction applied to simulated 40 nm diameter rings separated by 80 nm (a,b,c,g,h) or 120 nm (d,e,f,i). A simulated/experimental PSF for the 80 nm/120 nm separated rings was used, along with simulated drift curves in both cases. 50,000 frames were generated, divided into 100 datasets of 500 frames each. To make the simulation more realistic, the simulated localizations were fit and thresholded after being drifted. (a,d) Drifted image. The emitters are color-coded by frame number. (b,e) The drift-corrected image produced by driftCorrectKNN using default settings. (c,f) *x,y* perspective of the drift-corrected image. (g,i) The difference between the estimated and simulated *x, y, z*-drifts as a function of the absolute frame number for the (g/i) 80 nm/120 nm separated rings. (h) Corresponding drift correction plot for the 80 nm separation example.

**Figure S9.**
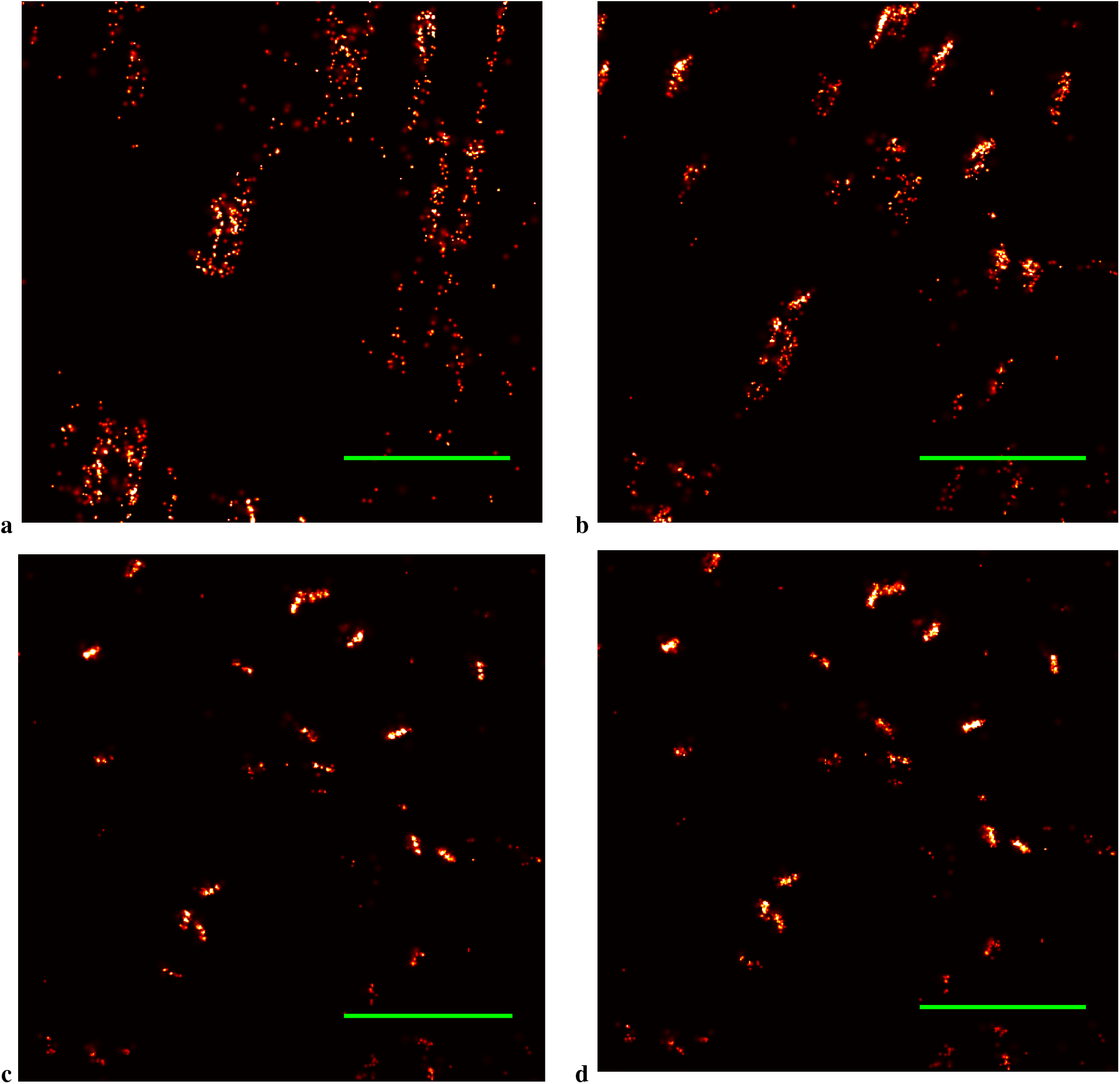
Example of internally reorganizing the datasets from an experiment examining ATTO655 20 nm nanorulers. The data was initially collected in a single dataset consisting of 15,000 frames. (a) Raw super-resolution image. (b) Post-processing drift correction was applied to one 15,000-frame dataset. (c) Post-processing drift correction was applied to 15 1,000-frame datasets. (d) Post-processing drift correction was applied to 150 100-frame datasets. All scale bars measure 1 *μ*m.

## Methods

### Frame connection

A blinking event often produces localizations across a sequence of frames. These localizations can be recognized as a single fluorophore and connected together to produce better precision. Frame connection takes the time-ordered localizations computed from a super-resolution dataset and attempts to combine them via a single emitter model in which the maximum distance and maximum frame gap between two localizations are specified (defaults are 1 pixel and 4 frames). The level of significance (LoS, default is 0.01) represents the minimum probability for which the null hypothesis that the two localizations come from a single emitter is not rejected. If the value computed for the probability is greater than the LoS and the other conditions hold, then the two localizations are combined. This process examines all localizations that satisfy the distance and frame gap constraints.

If two localizations have positions and localization errors (*x*_1_, *σ*_1_) and (*x*_2_, *σ*_2_), then the combined position and error is given by

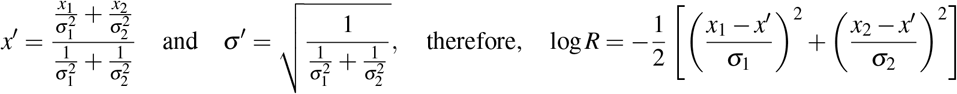

is the log-likelihood ratio (minus a constant term) that the two localizations represent a single emitter. The likelihood ratio is given by 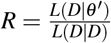, where the numerator is the likelihood *L* of the data *D* given the parameters *θ′* described above for the single emitter model, and the denominator is the likelihood of *D* given *D*, or one. *L* comes from the product of two Gaussians:

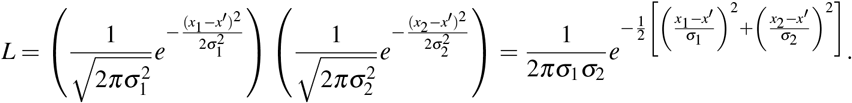

The corresponding *p*-value (which is compared to the LoS) is

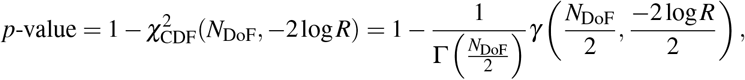

where *N*_DoF_ is the number of degrees of freedom and hence the spatial dimension, *N*_dim_, in the situation when two localizations are combined into one 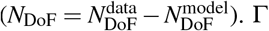 and *γ* are the gamma function and the lower incomplete gamma function, respectively. For *N*_DoF_ = 1,2,3,

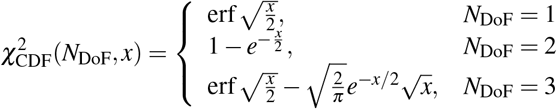

Note that in 2D (*N*_DoF_ = 2), the *p*-value is exactly *R*.

### Post-processing drift correction algorithm

MATLAB pseudocode drift correction algorithm operating on (*x, y*) or (*x,y, z*) localizations. The drift corrected coordinates, *X*, and the matrix of drift corrections indexed by dataset number and frame number, Δ*X*, are produced. *P*_degree_ is the degree of the polynomial fitting the intra-dataset drift correction. Note that the constant term is assumed to be zero. *N*_frames_ is the number of frames per dataset. Δ*X_i_* is the drift correction for each frame of the ith dataset. *p•,_j_T^j^* is the product of the 7^th^ column of p with the vector *T*, in which each component is raised to the *j*^th^ power. *X^c^* are drift corrected coordinates.

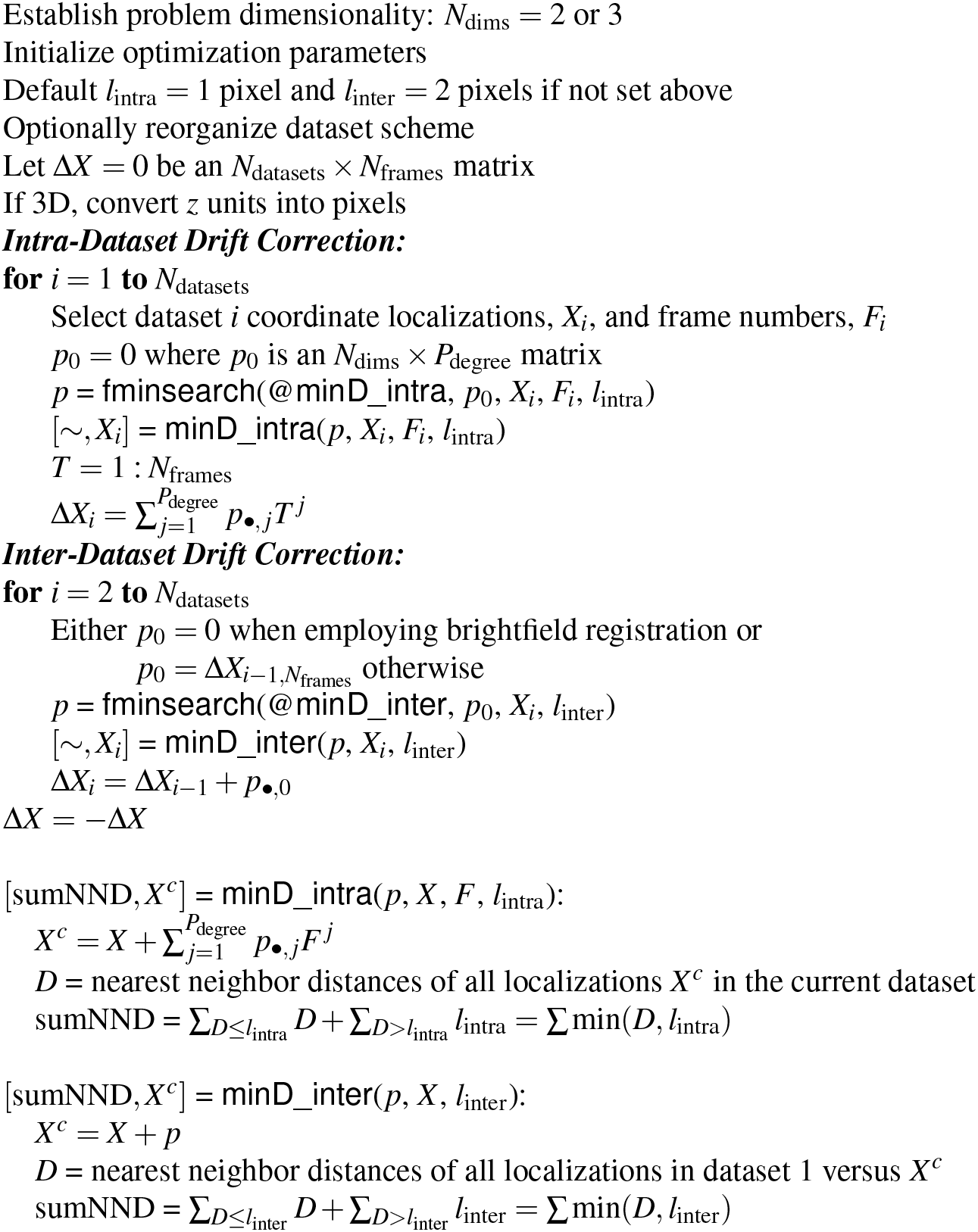

### RMSE analysis

The RMSE analysis, depicted in Supplementary Fig S6(a,b) and Supplementary Fig S7, plots the root-mean-square error (RMSE) between the true drift curve, 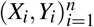, and the estimated drift curve, 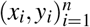, of simulated uniform randomly distributed single datasets of 2D emitters, versus the number of blinking event pairs in the datasets. The RMSE was computed by

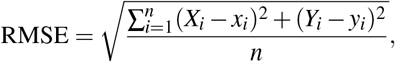

where a constant drift per frame was applied to the true locations to produce the drifted positions, which were then run through the intra-dataset portion of the post-processing drift correction algorithm. This definition of the RMSE considers the difference between the true drift curves at each time frame (as determined by the simulation) and the found (as determined by the intra-dataset portion of the post-processing drift correction algorithm).

## References

1. Snella, M. T. Drift Correction for Scanning-Electron Microscopy. Master’s thesis, Department of Electrical Engineering and Computer Science, Massachusetts Institute of Technology (2010).

2. Qiu, M. & Yang, G. Drift correction for fluorescence live cell imaging through correlated motion identification. In 2013 IEEE 10th International Symposium on Biomedical Imaging, DOI: 10.1109/ISBI.2013.6556509 (2013).

3. Mlodzianoski, M. J. et al. Sample drift correction in 3d fluorescence photoactivation localization microscopy. Opt. Express 19, 15009–15019, DOI: 10.1364/OE.19.015009 (2011).

4. Tang, Y., Wang, X., Zhang, X., Li, J. & Dai, L. Sub-nanometer drift correction for super-resolution imaging. Opt. Lett. 39, 5685–5688, DOI: 10.1364/OL.39.005685 (2014).

5. Han, R. et al. Drift-correction for single-molecule imaging by molecular constraint field, a distance minimum metric. BMC Biophys. 8, 1–14, DOI: 10.1186/s13628-014-0015-1 (2015).

6. Marturi, N., Dembélé, S. & Piat, N. Fast image drift compensation in scanning electron microscope using image registration. In IEEE International Conference on Automation Science and Engineering, CASE’13, 1–6, DOI: 10.1109/CoASE.2013.6653936 (2013).

7. Mantooth, B. A., Donhauser, Z. J., Kelly, K. F. & Weiss, P. S. Cross-correlation image tracking for drift correction and adsorbate analysis. Rev. Sci. Instruments 73, DOI: 10.1063/1.1427417 (2002).

8. Yothers, M. P., Browder, A. E. & Bumm, L. A. Real-space post-processing correction of thermal drift and piezoelectric actuator nonlinearities in scanning tunneling microscope images (2017). DOI: 10.1063/1.4974271.

9. Rahe, P., Bechstein, R. & Kühnle, A. Vertical and lateral drift corrections of scanning probe microscopy images. J. Vac. Sci. Technol. B 28, 4E31–C4E38, DOI: 10.1116/1.3360909 (2010).

10. Lidke, K. A., Rieger, B., Jovin, T. M. & Heintzmann, R. Superresolution by localization of quantum dots using blinking statistics. Opt. Express 13, 7052–7062, DOI: 10.1364/OPEX.13.007052 (2005).

11. Betzig, E. et al. Imaging intracellular fluorescent proteins at nanometer resolution. Science 313, 1642–1645, DOI: 10.1126/science.1127344 (2006).

12. Hell, S.W. Far-field optical nanoscopy. Science 316, 1153–1158, DOI: 10.1126/science.1137395 (2007).

13. Rust, M. J., Bates, M. & Zhuang, X. Sub-diffraction-limit imaging by stochastic optical reconstruction microscopy (storm). Nat. Methods 3, 793–796, DOI: 10.1038/nmeth929 (2006).

14. Lee, S. H. et al. Using fixed fiduciary markers for stage drift correction. Opt. Express 20, 12177–12183, DOI: 10.1364/OE.20.012177 (2012).

15. Schindelin, J., Rueden, C. T., Hiner, M. C. & Eliceiri, K. W. The imagej ecosystem: an open platform for biomedical image analysis. Mol. Reproduction Dev. 82, 518–529, DOI: 10.1002/mrd.22489 (2015).

16. Lombardot, B. Manual Drift Correction Plugin (2016). Accessed: February 28, 2018.

17. Petersen, S. B., Thiagarajan, V., Coutinho, I., Gajula, G. P. & Neves-Petersen, M. T. Image processing for drift compensation in fluorescence microscopy. In Farkas, D. L., Nicolau, D. V. & Leif, R. C. (eds.) Imaging, Manipulation, and Analysis of Biomolecules, Cells, and Tissues XI, vol. 8587, 85871H of Proceedings of SPIE, DOI: 10.1117/12.2004273 (2013).

18. Coelho, S. et al. Ultraprecise single-molecule localization microscopy enables in situ distance measurements in intact cells. Sci. Adv. 6, DOI: 10.1126/sciadv.aay8271 (2020).

19. Fan, X. et al. Three dimensional drift control at nano-scale in single molecule localization microscopy. Opt. Express 28, 32750–32763, DOI: 10.1364/OE.404123 (2020).

20. Parslow, A., Cardona, A. & Bryson-Richardson, R. J. Sample drift correction following 4d confocal time-lapse imaging. J. Vis. Exp. 51086, DOI: 10.3791/51086 (2014).

21. Sugar, J. D., Cummings, A. W., Jacobs, B. W. & Robinson, D. B. A free matlab script for spatial drift correction. Microsc. Today 22, DOI: 10.1017/S1551929514000790 (2014).

22. Smirnov, M. S., Evans, P. R., Garrett, T. R., Yan, L. & Yasuda, R. Automated remote focusing, drift correction, and photostimulation to evaluate structural plasticity in dendritic spines. PLoS ONE DOI: 10.1371/journal.pone.0170586 (2017).

23. McGorty, R., Kamiyama, D. & Huang, B. Active microscope stabilization in three dimensions using image correlation. Opt. Nanoscopy 2, DOI: 10.1186/2192-2853-2-3 (2013).

24. Wang, Y. et al. Localization events-based sample drift correction for localization microscopy with redundant cross-correlation algorithm. Opt. Express 22, 15982–15991, DOI: 10.1364/OE.22.015982 (2014).

25. Cizmar, P., Vladár, A. E. & Postek, M. T. Real-time scanning charged-particle microscope image composition with correction of drift. Microsc. Microanal. 17, 302–308, DOI: 10.1017/S1431927610094250 (2011).

26. Elmokadem, A. & Yu, J. Optimal drift correction for superresolution localization microscopy with bayesian inference. Biophys. J. 109, 1772–1780, DOI: 10.1016/j.bpj.2015.09.017 (2015).

27. Fazel, M., Wester, M. J., Rieger, B., Jungmann, R. & Lidke, K. A. Sub-nanometer precision using bayesian grouping of localizations. bioRxiv DOI: 10.1101/752287 (2019). https://www.biorxiv.org/content/early/2019/08/30/752287.full.pdf.

28. Friedman, J. H., Bently, J. L. & Finkel, R. A. An algorithm for finding best matches in logarithmic expected time. ACM Transactions on Math. Sofw. 3, 209–226, DOI: 10.1145/355744.355745 (1977).

29. Nieuwenhuizen, R. P. J. et al. Measuring image resolution in optical nanoscopy. Nat. Methods 10, 557–562, DOI: 10.1038/nmeth.2448 (2013).

30. Pallikkuth, S. et al. Sequential super-resolution imaging using dna strand displacement. PLoS ONE 13, e0203291, DOI: 10.1371/journal.pone.0203291 (2018).

31. Pallikkuth, S. et al. A matlab-based instrument control package for fluorescence imaging. Biophys. J. 114, 532a, DOI: 10.1016/j.bpj.2017.11.2912 (2018).

32. Riedl, J. et al. Lifeact: a versatile marker to visualize f-actin. Nat. Methods 5, 605–607, DOI: 10.1038/nmeth.1220 (2008).

33. Mazloom-Farsibaf, H. et al. Comparing lifeact and phalloidin for super-resolution imaging of actin in fixed cells. PLoS ONE 16, e0246138, DOI: 10.1371/journal.pone.0246138 (2021).

34. Liu, S., Kromann, E. B., Krueger, W. D., Bewersdorf, J. & Lidke, K. A. Three dimensional single molecule localization using a phase retrieved pupil function. Opt. Express 21, 29462–29487, DOI: 10.1364/OE.21.029462 (2013).

35. Fazel, M. et al. Bayesian multiple emitter fitting using reversible jump markov chain monte carlo. Sci. Reports 9, 1–10, DOI: 10.1038/s41598-019-50232-x (2019).

